# Notch directs telencephalic development and controls neocortical neuron fate determination by regulating microRNA levels

**DOI:** 10.1101/2022.09.16.508220

**Authors:** Jisoo S. Han, Elizabeth Fishman-Williams, Steven C. Decker, Keiko Hino, Raenier V. Reyes, Nadean L. Brown, Sergi Simó, Anna La Torre

## Abstract

The central nervous system (CNS) contains myriads of different types of cells produced from multipotent neural progenitors. Neural progenitors acquire distinct cell identities depending on their spatial position, but they are also influenced by temporal cues to give rise to different cell populations over time. For instance, the progenitors of the cerebral neocortex generate different populations of excitatory projection neurons following a well-known sequence. The Notch signaling pathway plays crucial roles this process but the molecular mechanisms by which Notch impacts progenitor fate decisions have not been fully resolved. Here, we show that Notch signaling is essential for neocortical and hippocampal morphogenesis, and for the development of the corpus callosum and choroid plexus. Our data also indicate that, in the neocortex, Notch controls projection neuron fate determination through the regulation of two microRNA (miRNA) clusters that include let-7, miR-99a/100, and miR-125b. Our findings collectively suggest that balanced Notch signaling is crucial for telencephalic development and that the interplay between Notch and miRNAs is critical to control neocortical progenitor behaviors and neuron cell fate decisions.

## INTRODUCTION

The mammalian telencephalon contains an unparalleled diversity of neural populations generated during development in a tightly regulated series of events. Despite the astounding intricacies of the mature cerebrum, the telencephalon arises from a relatively simple neuroepithelial sheet composed solely of neural progenitors (Rakic, 1972; Ramón y Cajal, 1899). An exquisitely orchestrated interplay of intrinsic and extrinsic factors choreographs the emergence of different territories along the antero-posterior, dorsal-ventral, and lateral-medial axes. The posterior medio-dorsal region will develop into the hippocampus, the cortical hem, and the choroid plexus, while the embryonic dorsal telencephalon will develop into the neocortex in the anterior and lateral aspects (Hebert & Fishell, 2008; Rallu *et al*, 2002; Wilson & Rubenstein, 2000).

At early stages of development, neural progenitors called radial glial cells (RGs) expand the whole thickness of the neocortex from the ventricular (apical) surface to the pial (basal) surface. As development proceeds and the cortex grows, the somas of the RGs remain close to lateral ventricles forming the ventricular zone where RGs can divide symmetrically to self-renew or asymmetrically to yield intermediate progenitors and post-mitotic neurons (Sidman & Rakic, 1973; Villalba *et al*, 2021). Importantly, these progenitors produce excitatory projection neurons in a conserved sequential manner (Frantz & McConnell, 1996; Luskin *et al*, 1988; Reid *et al*, 1995; Walsh & Cepko, 1988): around embryonic day (E) 10.5 in mice, the first post-mitotic neurons are generated and migrate away from the ventricular zone to form the preplate. Subsequent cohorts of neurons, split the preplate into marginal zone and subplate, establishing the cortical plate. Throughout the rest of neurogenesis, newly-born excitatory projection use the RGs as a scaffold to migrate radially through the existing layers and position themselves atop, forming an ‘inside-out’ lamination pattern (Noctor *et al*, 2004). Accordingly, the deeper neocortical layers (*e*.*g*., layer VI) are formed by early-born neurons, while the more superficial layers (*e*.*g*., layer II-III) contain late-born cells.

The Notch signaling pathway is a pivotal regulator of numerous developmental processes in the telencephalon, including regulating the balance between proliferation and differentiation of progenitor populations, cell fate acquisition, and glial cell specification, among other roles (Artavanis-Tsakonas *et al*, 1999; Dorsky *et al*, 1995; Irvine, 1999; Jadhav *et al*, 2006; Yoon & Gaiano, 2005). Ligands such as Delta-like or Jagged/Serrate bind to the transmembrane Notch receptors (Notch 1-4), causing in the proteolytic release of the Notch intracellular domain (NICD). NICD then translocates to the nucleus (Schroeter *et al*, 1998) and binds to a complex that includes RBPJ (recombination signal-binding protein for immunoglobulin κ J region, also known as CSL and CBF1), MAML1 (mastermind-like transcriptional co-activator1), p300, and other proteins to transcriptionally activate downstream genes (Fortini & Artavanis-Tsakonas, 1994; Smoller *et al*, 1990). Well-known effector targets of the Notch pathway include the HES (Hairy and Enhancer of Split) and HEY (Hairy Ears, Y-linked) families. Previous studies have reported that HES1-deficient mice exhibited accelerated neuronal differentiation in the neocortex while either HES1 (Ohtsuka & Kageyama, 2021) or HES5 (Bansod *et al*, 2017) overexpression led to an expansion of the neural progenitor pool and prolonged production of superficial layer neurons and astrocytes. Such findings suggest that the timing and levels of Notch signaling must be properly regulated to maintain the temporal control of neurogenesis.

Importantly, the Notch signaling pathway also engages in complex feedback loops with several microRNA (miRNAs) (Fishman *et al*, 2022). MiRNAs post-transcriptionally regulate the expression of Notch pathway components, including HES1 and HES5 (Georgi & Reh, 2011; Patterson *et al*, 2014). At the same time, Notch activity regulates the transcription of several miRNAs in different paradigms (Roese-Koerner *et al*, 2016; Singh *et al*, 2020; Tan *et al*, 2012). MiRNAs have recently emerged as key regulators of cortical fate acquisition and developmental timing and, in particular, let-7 and miR-125b are part of the heterochronic pathway that regulates many developmental transitions in bilaterally symmetrical animals and are key components of the molecular machinery that allows neural progenitors to generate late cell populations in the cortex and retina (La Torre *et al*, 2013; Shu *et al*, 2019).

Here, we show that balanced Notch signaling is necessary for proper telencephalic development, including the development of the neocortex, corpus callosum, hippocampus, and choroid plexus. Additionally, we show that Notch signaling regulates neurogenesis and the laminar organization of the cortex. At a molecular level, Notch coordinates the expression of several transcription factors, including bHLH neurogenic transcription factors as well as two miRNA clusters, *miR99ahg* and *miR100hg*, the host genes for the miRNAs miR-99a, let-7c, and miR-125b-2; and miR-100, let-7a, and miR-125b-1, respectively. Strikingly, we demonstrate that inhibition of these miRNAs partially rescues Notch gain-of-function phenotypes *in vivo*. Together our data indicate that complex interactions between the Notch pathway and miRNAs are essential for proper cell fate specification and overall telencephalic development.

## RESULTS

### Notch signaling regulates corpus callosum and hippocampal development

To investigate the roles of the Notch pathway during early telencephalic development, we generated Notch gain-of-function (GOF) and dominant negative (DN) mouse transgenic lines. A GOF strain was generated by crossing ROSA26^loxP-stop-loxP-Notch1-ICD^ (Murtaugh *et al*, 2003) with the Emx1-Cre driver (Gorski *et al*, 2002). The resulting mouse line, hereafter referred to as NICD, overexpresses Notch1-ICD in the dorsal telencephalon from embryonic day 10.5 (E10.5). Similarly, we generated a DN line by overexpressing a truncated MAML1 protein that acts as a dominant negative (ROSA26^loxP-stop-loxP-dnMAML1^ (Tu *et al*, 2005)) using the same Emx1-Cre driver (hereafter dnMAML). In both lines, CRE recombinase mediates excision of the loxP-flanked STOP cassette, allowing for the expression of either NICD or dnMAML and littermates containing no CRE were used as controls for all the experiments. Furthermore, in both cases, the constructs are inserted in the *ROSA26* locus and thus, they are equivalent, avoiding expression differences due to the surrounding genomic DNA.

Upon NICD overexpression, mice fail to thrive (Supplementary figure 1A) and die around two weeks of age whereas dnMAML mice show no differences in animal size or survival rates compared to their control littermates. At a global level, neither of these strains show significant brain size differences at birth (Postnatal day 0, P0), but by P10 dnMAML animals exhibit smaller telencephalons compared to the controls (Supplementary figure 1B).

Analyses of these mice at P0 revealed several telencephalic gross morphological defects. NICD brains exhibit enlarged lateral ventricles, aberrant hippocampi, thinner cortices, and agenesis of corpus callosum (Figure 1A-B, D-E, G-H, J-L). In contrast, dnMAML show smaller lateral ventricle volumes, smaller hippocampi, and dysgenesis of corpus callosum (Figure 1C, F, I-L), with the posterior aspect of the corpus callosum exhibiting stronger deficiencies, including misrouted axons (Figure 1F).

**Figure 1.**
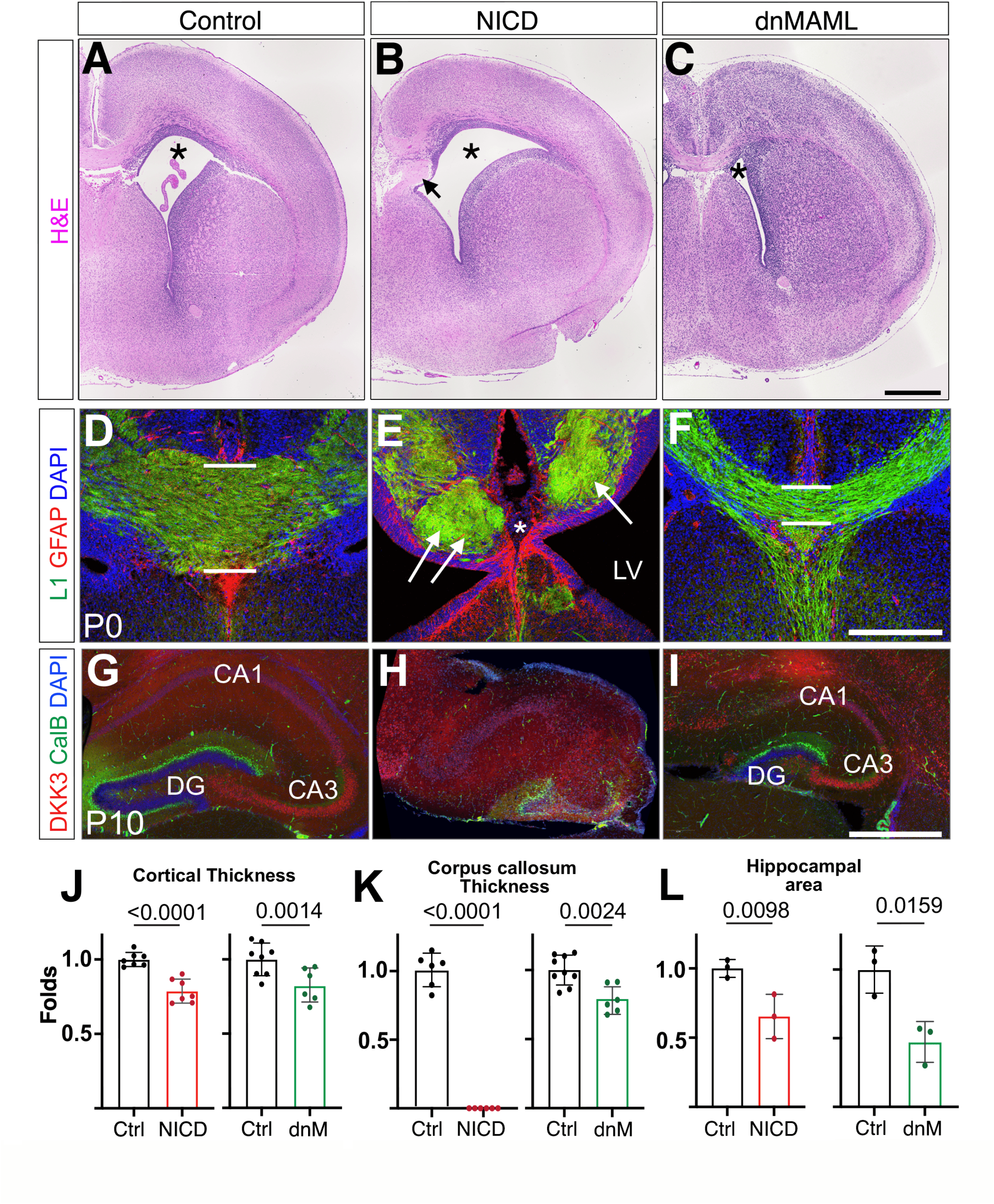
Morphological defects in NICD and dnMAML models. **A-C**. Hematoxylin and eosin staining (H&E) of Control (A), NICD (B) and dnMAML (C) P0 brains. Lateral ventricles are indicated with an asterisk. Arrow in B indicates axon Probst bundles. **D-F**. P0 cortical slices were immunolabeled against GFAP (red) and L1 (green) and counterstained with DAPI (blue). The thickness of the corpus callosum is indicated in D and F with white bars. White arrows in E point at Probst bundles and asterisk indicate lack of corpus callosum. **G-I**. P10 slices were immunolabeled against DKK3 (red) and Calbindin (green) and counterstained with DAPI (blue). Note the disorganization of the hippocampus in NICD (H). **J-L**. Quantifications of cortical thickness (J), corpus callosum thickness (K), and hippocampal area (L). Mean ± SEM. P-values were obtained using Student’s T-test. LV: lateral ventricle, DG: dentate gyrus, CA1-CA3: cornu ammonis hippocampal regions. Scale bars: 500μm A-C and G-I, 250μm D-F.

The corpus callosum is a large commissure that connects the right and left hemispheres and is formed by the axons of SATB2+ neurons that are found in all cortex layers, but are particularly abundant in layers II through IV (Britanova *et al*, 2008; Dobreva *et al*, 2006). The SATB2+ callosal neurons cross the midline through cell autonomous and non-cell autonomous mechanisms (Alcamo *et al*, 2008).

Since Notch signaling is known to regulate neuronal differentiation, one possibility is that the SATB2+ neurons perhaps are not correctly produced in NICD brains, thus resulting in a lack of callosal cells (Alcamo *et al*., 2008; Srivatsa *et al*, 2014). To test this hypothesis, we quantified the number of SATB2+ neurons in the neocortex of NICD mice at P0. Strikingly, NICD exhibits increased numbers of SATB2+ neurons compared to their littermate controls (2.49-fold increase, p-value: 0.004, Supplementary Figure 2A-B), indicating that the observed phenotype is not caused by a deficiency in the production of callosal neurons but possibly due to deficiencies in axon pathfinding or midline defects. In this direction, we observed Probst bundles by H&E staining and L1 axon immunolabeling that could correspond to aberrant bundles of axons that fail to extend through the midline (arrows in Figure 1B and E). To confirm this possibility, we labeled newly-born neurons with mCherry fluorescent protein in control and NICD embryos by *in utero* electroporation (IUE) at E13.5, corresponding with the age when the first SATB2+ neurons are born (Ozaki & Wahlsten, 1998) and we collected the mice at P0 for analysis. In control mice, we detected mCherry-labeled cells extending axons through the midline, whereas in NICD mice some mCherry+ axons extended towards the midline but failed to cross to the other hemisphere resulting in an aberrant accumulation of axon fascicles (Supplementary Figure 2C). The hippocampus is also affected in both NICD and dnMAML mice (Figure 1 G-I). The hippocampal formation comprises the dentate gyrus (DG), which includes the granule cells (Calbindin+) located in the granule cell layer (GCL), and the cornu ammonis (CA or hippocampus proper), which contains pyramidal neurons (DKK3+) (Han *et al*, 2020; Khalaf-Nazzal & Francis, 2013). NICD hippocampi are smaller than the controls and show a severe disorganization of both cell types at P10 without increased apoptosis at P0 (Figure 1H L, Supplementary figure 3). Conversely, upon dnMAML expression, the hippocampus shows seemingly normal organization of both DG and CAs, but all hippocampal regions are drastically reduced in size (Figure 1I, L). We also noticed varying degrees of mispositioning of DKK3+ neurons in the dnMAML hippocampi and in some cases, DKK3+ cells cross the upper DG blade and ectopically cluster in the hippocampal fissure (Supplementary figure 3D, arrows).

### Notch signaling is not a main mediator of dorsal telencephalic patterning but regulates Cajal-Retzius cell production

During the course of the CNS patterning, the dorsal telencephalic midline gets organized into three distinct regions: the choroid plexus (ChP), a small intermediate epithelium known as the cortical hem (CH), and the hippocampal primordium, which is contiguous with the neocortex (Subramanian & Tole, 2009) (Figure 2A-A’). The ChP is a nonneural epithelium responsible for producing the cerebrospinal fluid that fills the ventricles. The CH is as a classic developmental organizer as Wnt and BMP proteins secreted from this region play essential roles in patterning the adjacent regions (Furuta *et al*, 1997; Grove *et al*, 1998).

**Figure 2.**
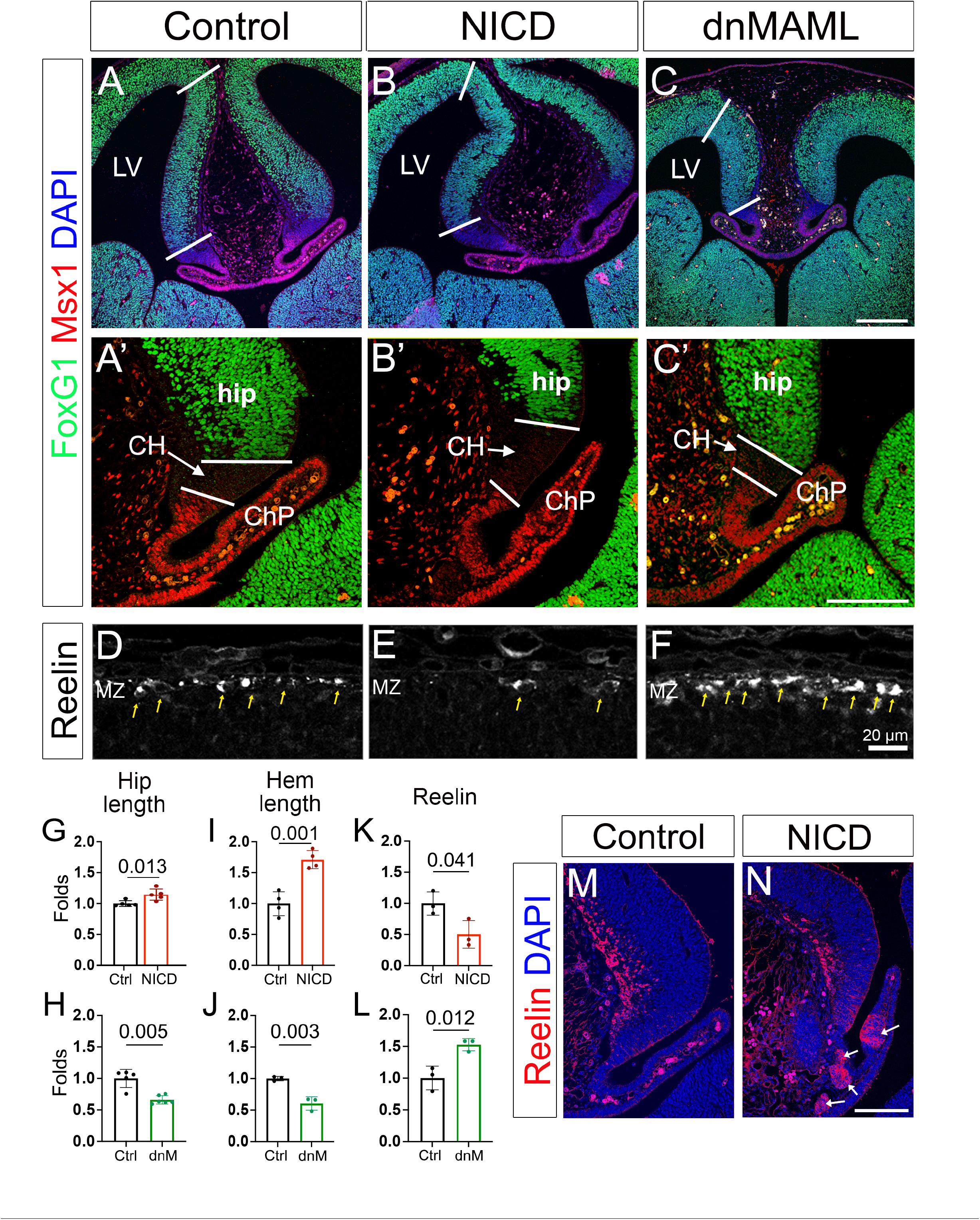
Midline patterning and production of Cajal-Retzius cells in NICD and dnMAML telencephalons. **A-C’**. Embryonic day 13.5 whole head sections were immunolabeled against MSX1 (red), FoxG1 (green), and counterstained with DAPI (blue). White bars indicate hippocampal primordia (A-C) and cortical hem (A’-C’) regions. **D-F**. Reelin (white) labeling in the marginal zone of the neocortex identifies Cajal-Retzius cells (yellow arrows). **G-L**. Quantifications of hippocampal length (G-H), cortical hem length (I-J), and number of Reelin+ cells/area (K-L) are shown as fold change compared to each control littermate. Mean ± SEM. P-values were obtained using Student’s T-test. **M-N**. Reelin stainings reveal the presence of ectopic patches of Cajal-Retzius cells in the choroid plexus of NICD mice at E13.5 (white arrows). LV: lateral ventricle;, hip: hippocampal primordia; CH: cortical hem; ChP: choroid plexus; MZ: marginal zone. Scale bars: 250μm A-C, 100μm A’-C’ and M-N, 20μm D-F.

Since both the NICD and dnMAML mice exhibit abnormal hippocampi and enlarged or smaller lateral ventricles respectively, we hypothesized that the dorsal telencephalic midline patterning could be affected in our models. Supporting this idea, a triple knockout of the Notch signaling effectors *Hes1, Hes3*, and *Hes5* exhibited defects in ChP development(Imayoshi *et al*, 2008). To identify possible patterning alterations, we labeled E13.5 whole-head coronal sections with FOXG1 and MSX1 to distinguish the hippocampal primordium, CH, and ChP regions (Figure 2A-C’). NICD mice exhibit elongated hippocampal primordia and CH, while these structures are significantly shorter in the dnMAML compared to their littermate controls (Figure 2A-C’, G-J). These changes could be reflecting alterations in cell cycle dynamics, but the general patterning and localization of these territories is maintained.

An important function of the CH is the production of Cajal-Retzius cells that secrete the extracellular glycoprotein Reelin (Soriano & Del Rio, 2005), which is essential for cortical and hippocampal neuron migration and lamination (Borrell *et al*, 2007; Sekine *et al*, 2011). Since the size of the CH is altered in both NICD and dnMAML mice, we labeled and quantified the number of Cajal-Retzius cells, using Reelin as a marker. Although the CH is larger in NICD brains, we found a reduction in Reelin+ cells in both the cortical marginal zone (Figure 2D, E, K, Supplementary 4A, C) and in the hippocampus at P0 (Supplementary Figure 4B). Interestingly, despite fewer Reelin+ cells are present in the cortex and hippocampus in NICD mice, we identified ectopic patches of Reelin+ cells within the ChP (Figure 2 M-N). Reciprocally, dnMAML mice showed a significant increase in Reelin+ cells at E13.5 in the cortical marginal zone (Figure 2C, L), although no changes were observed in the hippocampus (Supplementary Figure 4D).

### Early-born projection neuron production is limited by Notch signaling during neocortical development

Since our NICD animals showed increased numbers of SATB2+ cells, we assessed if other cortical cell types were also affected by NICD overexpression. To avoid staining or counting bias, we used a semi-automatic cell counter platform (RapID (Sekar *et al*, 2021)) and we normalized each quantification to their corresponding littermate controls. In NICD cortices, we observed a dramatic reduction of CTIP2+ and TBR1+ projections neurons and an increase in CUX1+ cells in comparison to control brains (Figure 3A-D). Despite the significant changes in the ratio of cell populations in these cortices and the reduction in Reelin+ cells, lamination was largely normal, with CUX1+ projection neurons located at the top of the cortex and TBR1+ neurons located in the most apical layer of the cortical plate (Figure 3A).

**Figure 3.**
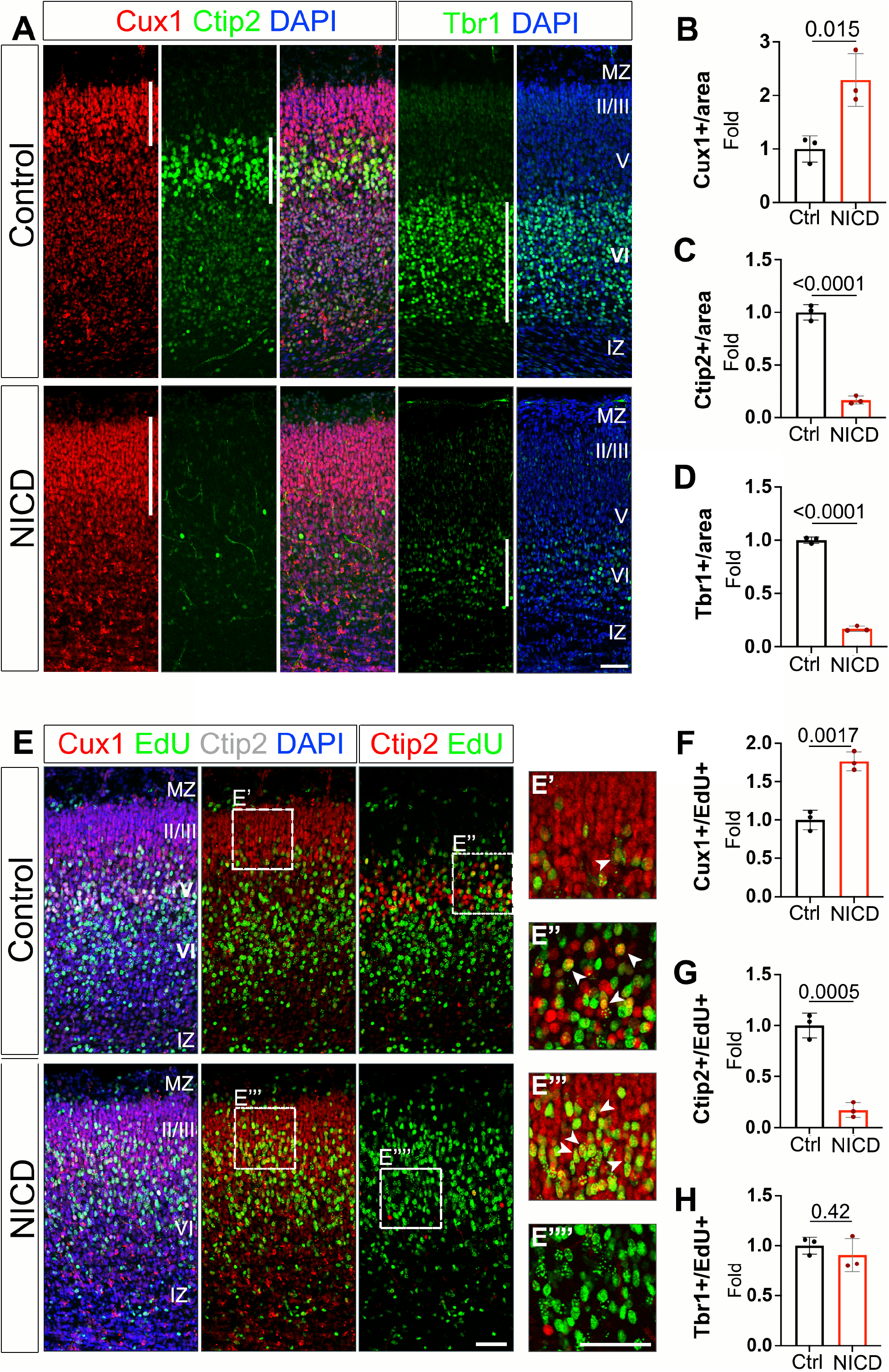
NICD neocortices exhibit increased ratios of upper-layer neurons. **A**. P0 cortical brain section immunolabeled with CUX1 (red), CTIP2 (green), and TBR1 (green) antibodies, and counterstained with DAPI (blue). **B-D**. Quantifications of the number of cells per area are shown as fold change compared to their control littermates. **E**. Cortical section labelings of EdU (green), CUX1 (red), CTIP2 (white left, red right) are counterstained with DAPI (blue). **F-H**. Quantifications of the number of cells per area are shown normalized to their corresponding control littermates. Mean ± SEM (B-D and F-H). P-values were obtained using Student’s T-test. Scale bars: 50μm.

To further investigate the changes in cell subpopulations, we performed birth-dating experiments using EdU (5-ethynyl-2’-deoxyuridine) to label dividing progenitors. We injected EdU at E13.5 and then we analyzed the cortices at P0 to assess the fate outcomes of the EdU-labeled progenitors. The number of neurons that were both EdU+ and CUX1+, CTIP2+, or TBR1+ were quantified and normalized by the total number of EdU+ neurons (Figure 3E-H). Whereas in control brains EdU-labeled E13.5 neural progenitor cells mostly gave rise to CTIP2+ layer V neurons (Figure 3E, G), we observed a strong decrease of EdU+ CTIP2+ cells in NICD brains. Concomitantly, the presence of EdU+ CUX1+ cells increased almost two folds in NICD brain in comparison to their control littermates (Figure 3E, F). At this stage of development, RGs have already passed beyond the period of production of layer VI neurons, and we did not observe a significant change in TBR1+ cell production (Figure 3H). Next, we evaluated the consequences of blocking Notch signaling in the developing cortex using the dnMAML model. Unfortunately, the anti-CUX1 antibody that we were using (Santa Cruz Biotechnology) was discontinued and we had to switch to a new vendor (Proteintech). The new antibody only labels CUX1+ neurons efficiently after P10 and thus, we switched the age of all subsequent CUX1 analyses from P0 to P10. In this case, we found a significant decrease in CUX1+ projection neurons, whereas CTIP2+ and TBR1+ neurons were overrepresented (Figure 4A-G). We also performed birth-dating experiments to measure whether the changes in cortical neuron composition are linked to changes in the timing of neurogenesis. Similar to the experiments described before, we injected EdU to pregnant females at E13.5 and collected the pups at birth or P10 for analysis. We found a significant decrease in EdU+ CUX1+ neurons and a corresponding increase in EdU+ CTIP2+ and EdU+ TBR1+ neurons, suggesting that the neural progenitors in the dnMAML mice continue to produce TBR1+ neurons beyond the normal time window of layer VI neurogenesis (Figure 4J-O).

**Figure 4.**
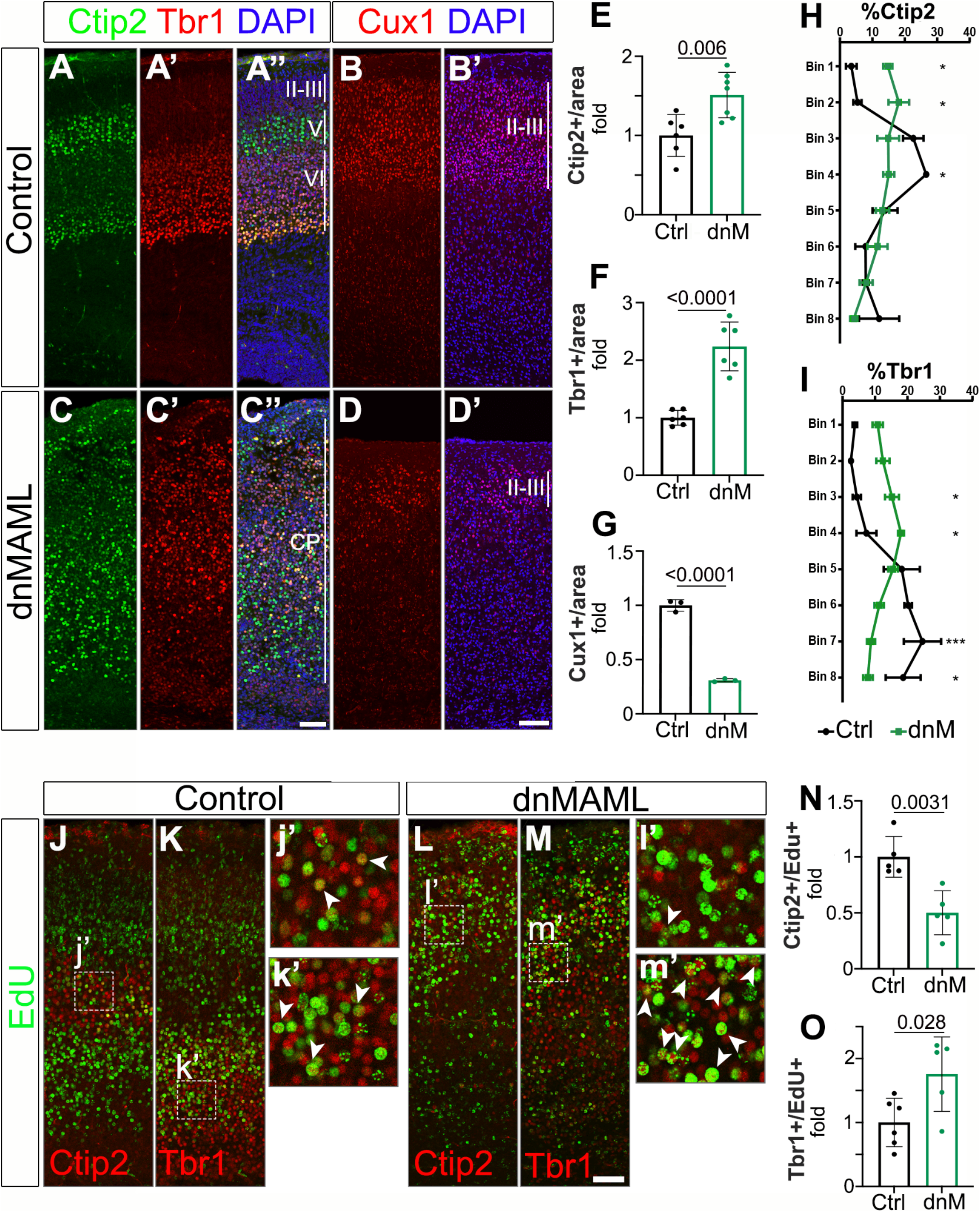
dnMAML neocortices exhibit increased numbers of deep-layer neurons and lamination defects. **A-C’**. P0 cortical coronal section immunolabeled with CTIP2 (green) and TBR1 (red) (A-A’’ and C-C’’) and P10 coronal section immunolabeled with CUX1 (red) antibodies. Tissues were counterstained with DAPI (blue). **E-G**. Quantifications of the number of cells per area are shown as fold change compared to their control littermates. **H-I**. Distribution of Ctip2+ (H) and Tbr1+ (I) cells in control (Ctrl) and dnMAML (dnM) in P0 cortical brain slices. **J-M’**. Cortical section labelings of EdU (green), CTIP2 (red), and TBR1 (red) are counterstained with DAPI (blue). **N-O**. Quantifications of the number of cells per area are shown normalized to their corresponding control littermates. Mean ± SEM. P-values in (E-G) and (N-O) were obtained using Student’s T-test. For cell distribution in (H-I), multiple unpaired T tests (one per bin) with Welch correction were performed (*, Adjusted P value <0.05; ***, Adjusted P value <0.001). Scale bar: 50μm.

In contrast with the normal lamination observed in Notch GOF, dnMAML mice neocortices also show severe disruption of the cortical layers in the dorsomedial region with milder lamination defects in the lateral aspects of the neocortex (Supplementary Figure 5). To quantify this phenotype, we divided the cortical plate into eight bins and counted the number of CTIP2+ and TBR1+ neurons in each of the bins. In control animals, TBR1+ neurons are mainly positioned at the bottom of the cortical plate as expected (bins 6-8 contain 63.5% of all TBR1+ neurons) and CTIP2+ cells are enriched in more basal locations (bins 3-4 contain 49.2% of all CTIP2+ cells). Conversely, both CTIP2+ and TBR1+ neurons were found dispersed across the whole thickness of the cortical plate in the dnMAML samples (bins 6-8 contain only 27.8% of all TBR1+ cells while bins 3-4 include 29.9% of CTIP2+ neurons) (Figure 4H-I).

Despite MAML having clear roles in the Notch signaling pathway, several studies suggest broader functions for MAML1 as a cofactor for multiple signaling pathways, including Wnt, Hippo, and Sonic Hedgehog (Alves-Guerra *et al*, 2007; Kim *et al*, 2020; Quaranta *et al*, 2017; Zema *et al*, 2020). To determine whether the defects observed in dnMAML mice are caused by its role in the MAML-RBPJ-NICD trimeric protein complex, we generated a Notch1 conditional knockout line by crossing Notch1^f/f^ mice (Yang *et al*, 2004) with the Emx1-CRE driver (Notch1cKO mice). Similar to dnMAML1 mice, Notch1cKO exhibited dysgenesis of the corpus callosum and smaller lateral ventricles and hippocampi (Supplementary Figure 6A-D). We also observed thinner cortices that had considerably reduced the upper layers (II-III) and dispersed cortical neurons, but this phenotype was not as severe as in the dnMAML model (Supplementary Figure 6E, F). We performed birth-dating experiments as described above. These experiments showed a 2.4-fold increase in TBR1+ population production (*i*.*e*., TBR1+ EdU+) compared to littermate controls, whereas no differences were observed for CTIP2+ population (Supplementary Figure 6G, H). These data indicate that Notch1cKO closely mimic dnMAML phenotypes, suggesting that the overall defects we observe in dnMAML mice are mainly due to the imbalance downstream of Notch.

Together, our data show that Notch signaling regulates neurogenesis and limits the production of early-born cell fates (TBR1+ and CTIP2+), while increasing the genesis of late-born cells (CUX1+).

### Notch signaling regulates radial glia cell cycle dynamics

Notch signaling is required for maintaining the progenitor pool and *Hes* genes downstream of Notch repress bHLH transcription factors, especially those with proneural functions (Dennis *et al*, 2019; Huang *et al*, 2014; Imayoshi *et al*, 2010; Jadhav *et al*., 2006; Kageyama *et al*, 2009; Maurer *et al*, 2014; Nieto *et al*, 2001). Similarly, overexpression of Delta1, HES1, or activated Notch1 prolongs mitotic activity in different types of progenitor cells (Austin *et al*, 1995; Del Bene *et al*, 2008; Dorsky *et al*., 1995; Kageyama *et al*., 2009). For these reasons, we characterized the cortical neural progenitors in our transgenic models at E13.5. We observed decreased numbers of TUJ1+ post-mitotic neurons at E13.5 in NICD mice as expected (Figure 5A, right panels), but interestingly, we also found that all NICD embryos exhibit a complete depletion of TBR2+ intermediate progenitors (Figure 5A). To further validate that the intermediate progenitors were absent as opposed to a downregulation of TBR2 protein, we labeled all mitotic cells with Phosphohistone H3 (PH3). While we did not observe any changes in the number of mitotic cells adjacent to the ventricle (RGs), the basally located PH3+ cells (intermediate progenitors) were absent in NICD mice (Figure 5B-D).

**Figure 5.**
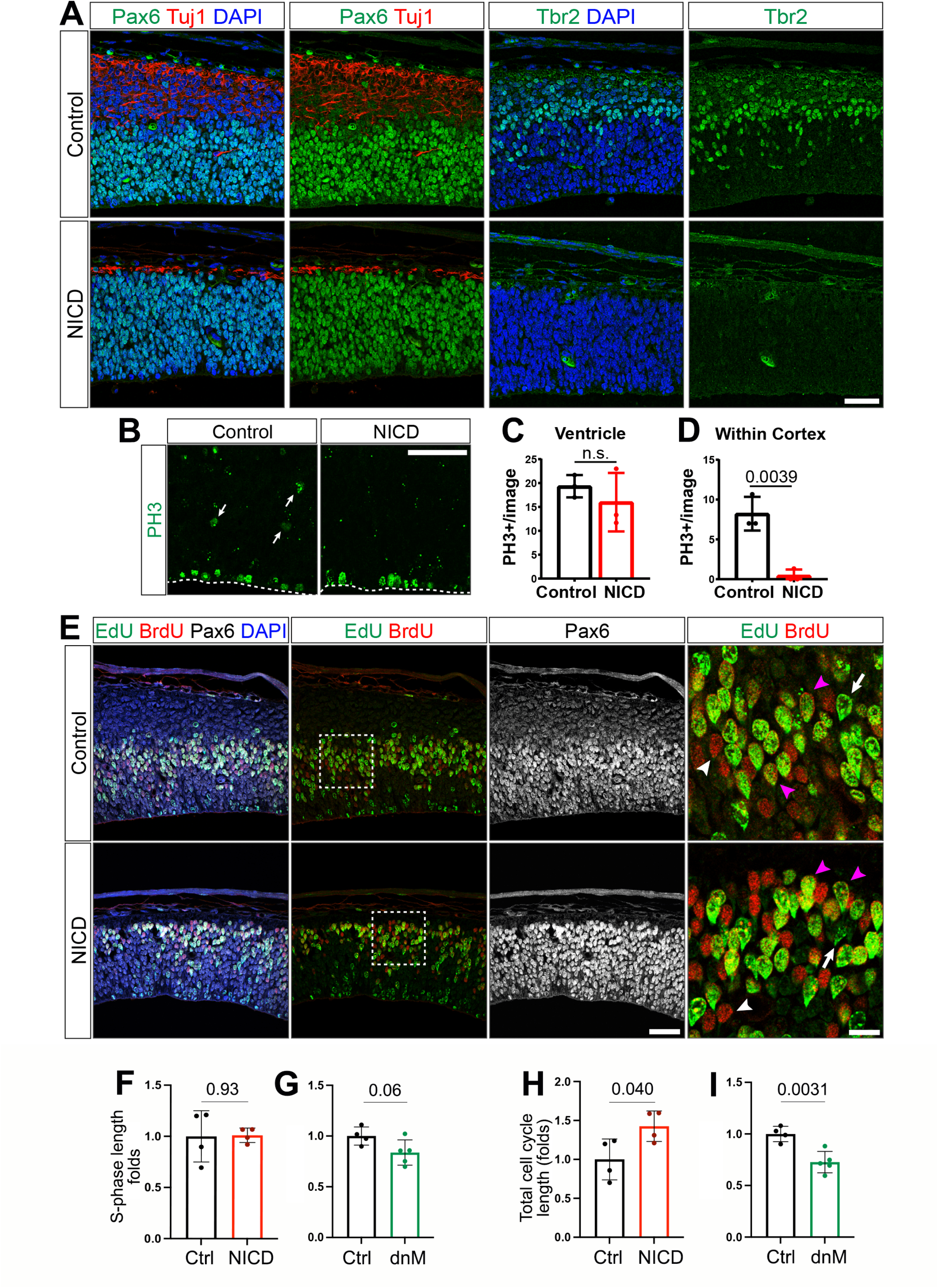
Notch regulates Radial Glia cell cycle dynamics. **A**. Cortical slices from E13.5 control and NICD embryos immunostained against Pax6 (green), ß-III-Tubulin (Tuj1, red), and Tbr2 (green) counterstained with DAPI (blue). **B**. Cortical slices from E13.5 control and NICD embryos immunostained against phosphor-Histone3 (PH3, green). **C-D**. Quantification of PH3+ cells in the ventricular surface (C) and anywhere else in the cortex area above the ventricular surface (D). **E**. E13.5 cortical section labelings of EdU (green), BrdU (red), and Pax6 (white). Panels on the left are also counterstained with DAPI (blue). **F-G**. Quantification of S-phase length in NICD (F) and dnMAML (dnM) (G) mice in comparison to their control littermates. **H-I**. Quantification of total cell cycle length in NICD (H) and dnMAML (dnM) (I) mice in comparison to their control littermates. Mean ± SEM (F-I). P-values were obtained using Student’s T-test. Scale bars: 50μm A, B and E (except for the right panels in E, 20μm).

To further characterize the cell cycle dynamics in our different genetic models, we measured the length of the cell cycle using dual-window labeling with the thymidine analogs EdU and BrdU (5-bromo-2’-deoxy-uridine) as described before(Harris *et al*, 2018). Pregnant mice at E13.5 were injected with a pulse of EdU followed by a pulse of BrdU two hours later. All animals were euthanized 30 minutes after the second pulse (150 minutes total) and the tissues were processed and stained for EdU, BrdU, and PAX6. Since NICD mice were missing the TBR2+ intermediate progenitors, we limited the quantification to PAX6+ apical RGs. RGs labeled by EdU but not BrdU (PAX6+ EdU+ BrdU-) left S-phase during the 2-hour period between pulses. The ratio of PAX6+ EdU+ BrdU-cells over the total number of cells in S-phase (PAX6+ EdU+) equals 2h/Time of S-phase (2h/Ts). The ratio between the number of cells in S phase at one given timepoint (PAX6+ BrdU+) and the total PAX6+ proliferating population is proportional to the ratio Ts/total cell cycle time (Ts/Tc). Using these parameters, we estimated the percentage of cells in S phase for each sample and then calculated the average Ts and Tc and we normalized each value to their littermate controls to avoid staining or imaging bias.

While the length of S-phase was not significantly altered in any of the models (9.02h in control, 9.12h in NICD and 9.20h in dnMAML Fig. 6E-G), the total length of cell cycle was significantly longer in NICD progenitors (7.9 hours longer or 1.42-fold increase ±0.19, p-value: 0.040) and shorter in dnMAML RGs (6.7 hours shorter or 1.38-fold reduction ±0.59, p-value: 0.003).

**Figure 6.**
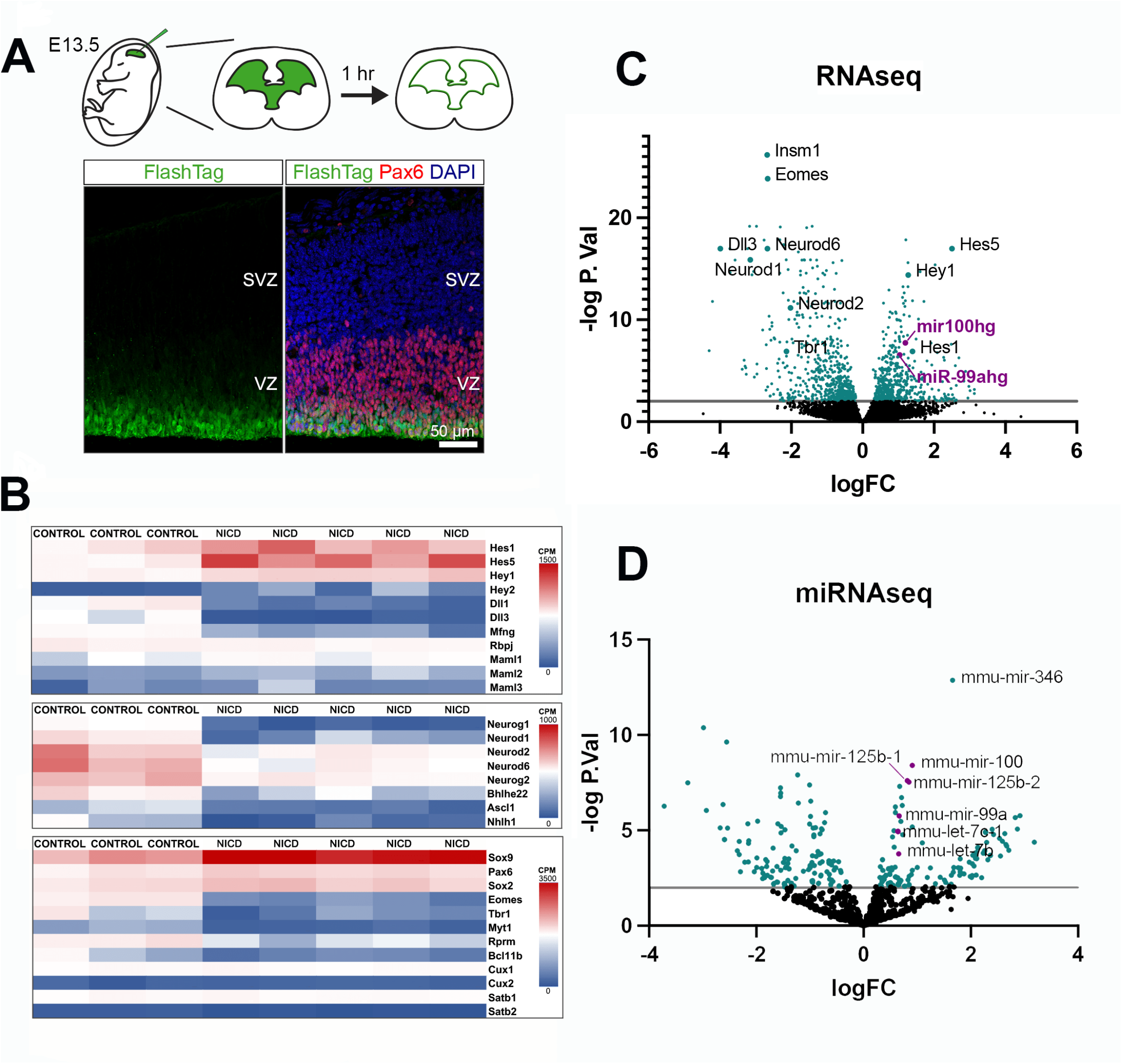
Transcriptional changes in NICD Radial Glia. **A**. E13.5 control and NICD embryos were injected with CSFEs (FlashTag) labeling cells surrounding the ventricles (top). In the cortex, only RGs, identified by the expression of Pax6 (red), closest to the ventricular surface were labeled with FlashTag (green) one hour post-injection (bottom). **B**. Heatmaps showing RNA-seq analysis of selected Notch signaling-related genes (top), bHLH genes (middle), and cortical markers (bottom). **C-D**. Volcano plots representing differential gene (C) and miRNA (D) expression between NICD (n=5) and control (n=3) RGs samples. Ratio of counts per million between NICD and control per gene or miRNA is plotted. The x-axis represents the logarithmic fold ration of NICD/control per gene and miRNA identified. The y-axis represents the logarithmic adjusted p-value (false discovery rate) calculated by the Benjamini-Hochberg Procedure. Genes with fewer than 5 counts per million reads in all samples were filtered prior to analysis, leaving 12,556 genes. MiRNAs present in fewer than 3 samples were filtered prior to analysis, leaving 779 miRNAs.

Taken together, these data show that activation of Notch signaling results in depletion of intermediate progenitors and lengthening of the cell cycle without affecting the total length of the S-phase.

### Overactivation of Notch signaling results in transcriptomic changes of Notch effectors, bHLH transcription factors, and miRNAs let-7, miR-99a/100, and miR-125b

To identify the downstream targets of Notch activation that may play a role in neurogenesis and/or cell cycle regulation within the progenitor population, we performed RNA sequencing (RNAseq) of neocortical RGs at E13.5. To that goal, we labeled RGs using FlashTag labeling (Govindan *et al*, 2018). As described previously, FlashTag utilizes carboxyfluorescein esters (CFSEs) that, when injected into the ventricles, label cells in contact with the cerebrospinal fluid. Since the apical RGs are transiently in contact with the ventricle walls during mitosis, this technique allows for a specific labeling of the RG population. We confirmed that 1 hour after injection, all FlashTag-labeled cells were PAX6+ RGs (Figure 6A). Next, we used FlashTag to label RGs in control (n=3) and NICD (n=5) littermate embryos at E13.5. We dissected the neocortices 1 hour post -injection, the labeled RGs were isolated using FACS, and we performed RNAseq. Multidimensional scaling analysis showed a clear clustering of all the NICD samples (Supplementary Figure 7A). Gene ontology (GO) enrichment analyses using PANTHER revealed that processes overrepresented in NICD cortices include the terms: “regulation of Notch signaling pathway” (GO:0008593, p-value: 0.00135), “negative regulation of cell differentiation” (GO:0045596, p-value 2.22×10^−9^), “cell fate commitment” (GO:0045165, p-value: 1.73×10^−8^), and “regulation of cell cycle” (GO:0051762, p-value: 0.00026); processes downregulated include the terms: “neuron differentiation” (GO: 0030182, p-value: 4.36×10^−17^), “forebrain development” (GO:0030900, p-value: 1.65×10^−10^), and “axon guidance” (GO:0007411, p-value: 0.00148)(Supplementary Figure 7B).

As expected, known Notch effectors such as *Hes1, Hes5, Hey1*, and *Hey2* were upregulated by NICD, while the Notch receptors *Dll1, Dll3* were downregulated as well as *Mfng* (Manic Fringe Homolog), a glycosyltransferase that modulates Notch activity (Figure 6B-C). *Hes* and *Hey* genes negatively regulate the expression of proneural basic helix-loop-helix (bHLH) transcription factors in several contexts and accordingly, we observed a reduction in *Neurog1, Neurog2, Neurod1, Neurod2, Neurod6*, and *Ascl1* (Figure 6B-C). Even though we only analyzed the transcriptome of the RGs, we observed a significant downregulation of some deep layer markers (*e*.*g*., *Tbr1, Myt1* and *Rprm*), Layer V (*Bcl11b/Ctip2*), and intermediate progenitor genes (*e*.*g*., *Eomes/Tbr2*), but we did not observed differences in upper layer markers (*e*.*g*., *Cux1* and *Satb2*).

Strikingly, we also observed upregulation of *miR100hg* and *miR99ahg* which are the host genes for two microRNA (miRNA) clusters (Figure 6C, purple data points). *MiR100hg* (miR-100 host gene) includes miR-100, let-7a-2, and miR-125b-1 while *miR99ahg* (mir-99a host gene) encodes for miR-99a, miR-125b-2, and let-7c. Upregulation of these miRNAs in NICD samples was further confirmed by miRNA sequencing of FlashTag-labeled purified RGs (Figure 6D and Supplementary Table 2), as described before.

### Let-7, miR-125b, and miR-99a/100 are required downstream of Notch to restrict early-born projection neuron fates

To test whether the upregulation of let-7, miR-125b and/or miR-99a/100 play any roles in the cortical phenotypes observed, we performed IUE using microRNA sponges (Barta *et al*, 2016; La Torre *et al*., 2013) to inhibit miRNA activity in NICD mice, and we analyzed the effects on cell fate. Plasmids expressing specific miRNA sponges together with an mScarlet plasmid were IUEd into NICD E13.5 embryos and the electroporated brains were collected at P0 (Figure 7A-F). These samples were processed, labeled with CTIP2 antibodies, and numbers of CTIP2+ mScarlet+ cells were determined. Whereas in control animals we observed 19.4% ± 8.4 of CTIP2+ mScarlet+/mScarlet+ cells, we did not observe any CTIP2+ mScarlet+ cells in NICD brains (Figure 7A-B), in agreement with our data showing that upon NICD overexpression, RGs generate upper-layer cells instead of layer V neurons at E13.5. Importantly, when let-7 was inhibited by means of a let-7 sponge, we observed a partial although non-significant rescue in the number of CTIP2+ electroporated neurons (5.56% ± 2.9, Figure 7C,G). In contrast, we did not observe any changes when miR-125b or miR-100 were inhibited (0% and 2.83% ± 2.6, respectively, Figure 7D,E,G). Notably, when we electroporated the three sponges together (let-7, miR-125b and miR-100) in NICD mice, significantly more electroporated cells were now CTIP2+ (11.15% ± 7.4, Figure 7F,G). Overall, these data suggest that these miRNAs are downstream effectors of Notch signaling and necessary to produce upper-layer neurons, possibly through epistatic mechanisms.

**Figure 7.**
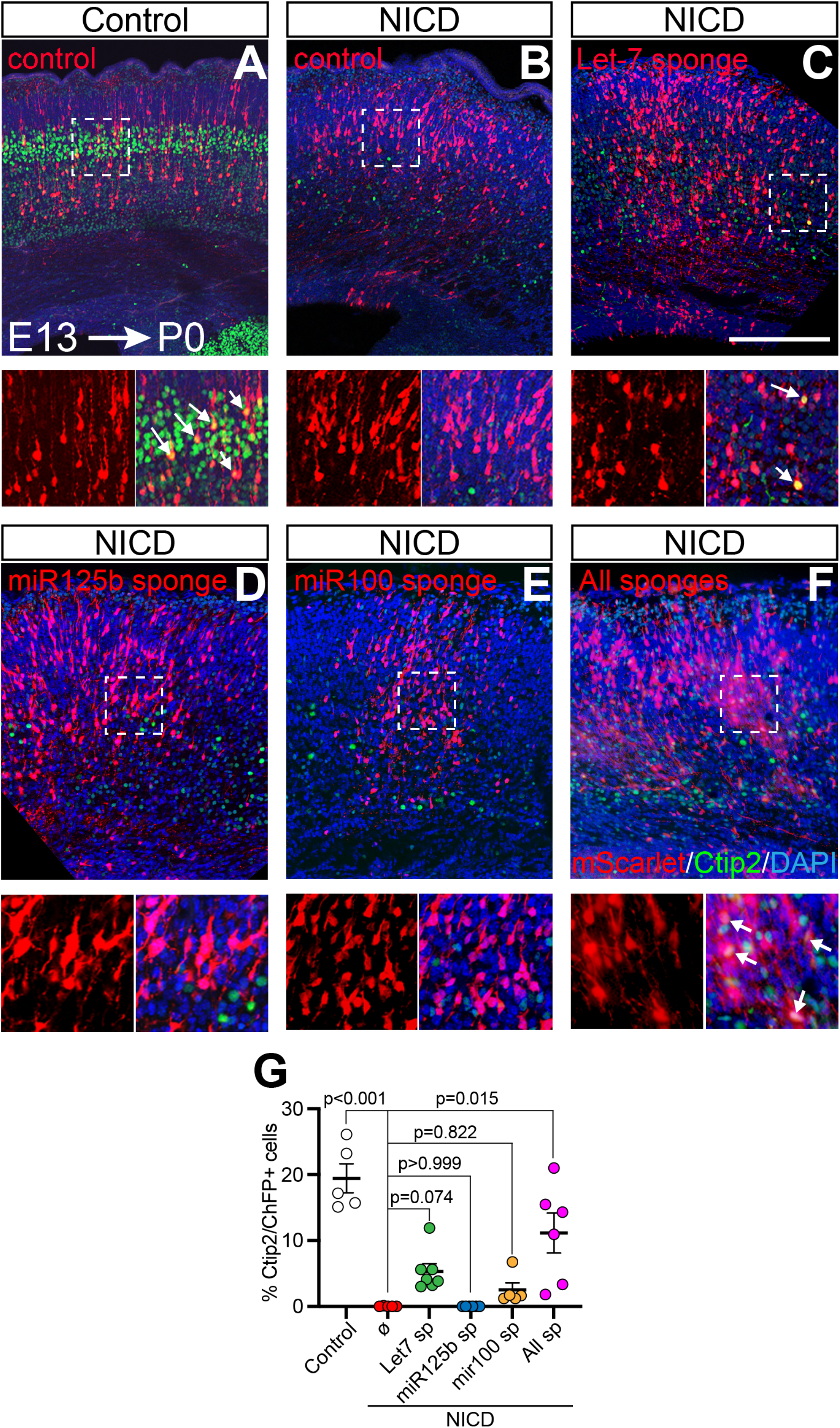
Inhibition of let-7, miR-125b, and miR-100 expression rescues cortical cell fate defects in NICD mice. **A-B**. Representative images of control (A) and NICD (B) mice electroporated with control plasmid (mScarlett, red) and NICD mice co-electroporated with mScarlett and miRNA sponges against let-7 (C) miR-125b (D), miR-100 (E), or all three sponges (F). Quantification of the percentage of Ctip2+/mScarlet+ cells in each condition. Mean ± SEM. Adjusted P values were obtained with Kruskal-Wallis test and Dunn’s post-hoc test. Scale bar: 250μm.

## DISCUSSION

### Multifaceted functions of Notch during telencephalic morphogenesis

Notch has long been recognized for its roles in cell specification, patterning, differentiation, and regeneration (Artavanis-Tsakonas *et al*., 1999; Bosze *et al*, 2020; Conner *et al*, 2014; Imayoshi *et al*., 2010; Jadhav *et al*., 2006; Maurer *et al*., 2014; Nelson *et al*, 2007; Riesenberg *et al*, 2009; Sestan *et al*, 1999). However, the extent of its contributions in telencephalic development remains unresolved. Utilizing GOF and DN transgenic mouse lines, we demonstrate that balanced Notch signaling is required for hippocampal, ChP, and corpus callosum development, and we also show that Notch is key to regulate neurogenesis in the neocortex.

Given the vast array of gross morphological defects observed in both GOF and DN models, we examined whether Notch regulates the patterning of the dorsal telencephalic midline, perhaps affecting the development of the hippocampus and ChP, changing brain fluid homeostasis and the volume of the ventricles. Although there are obvious differences in the size of the hippocampal, CH, and ChP areas in the transgenic mice, the establishment of the different territories is not affected by Notch changes. However, while the ChP is properly patterned and establishes a sharp boundary with the CH, we identified ectopic Reelin+ cells in the ChP region at E13.5 in our NICD model. A previous study using lineage-tracing analysis of the prospective ChP region indicated that these progenitors sequentially give rise to Reelin+ Cajal-Retzius cells first and later to nonneural ChP epithelial fates (Imayoshi *et al*., 2008). Inactivation of *Hes1, Hes3* and *Hes5* genes led to an enhanced development of Cajal-Retzius cells at the expense of ChP cell fates, suggesting that Notch signaling regulates the progenitor transition from cells that produce Cajal-Retzius cells to progenitors that will generate nonneural ChP epithelium cell identities. However, around E12.5, the levels of HES1 and HES5 are downregulated after the ChP cell fate is specified (Imayoshi *et al*., 2008). The presence of ectopic Cajal-Retzius cells in our data suggests that this later dowregulation of Notch signaling is required to maintain the ChP fate and that ChP progenitors can transdifferentiate to neural fates upon sustained HES1/5 activity. The ectopic Cajal-Retzius cells observed could alternatively be the consequence of aberrant migratory patterns from the CH. In fact, Notch has been shown to regulate migration patterns in the cortex through interactions with the Reelin-Dab1 signaling pathway (Hashimoto-Torii *et al*, 2008). In the cortex, a large reduction in Reelin+ Cajal-Retzius cells (50% reduction by E13.5 and 60% by P0) does not affect cell lamination in NICD mice, in agreement with other studies in which the ablation of the CH, leading to a severe reduction in Cajal-Retzius cells, did not affect the neocortical layer organization (Yoshida *et al*, 2006). On the other hand, dnMAML mice show severe cortical lamination defects. This phenotype resembles the defects observed in Emx1-Cre; RBPJ^f/f^ mice, in which alterations in the structure of the RGs were also observed (Son *et al*, 2020).

### Balanced Notch signaling is essential for the development of the corpus callosum

Transgenic models overexpressing *Hes5* in the neocortex (Bansod *et al*., 2017) and *Hes1/Hes3/Hes5* and *RBPJ* knockouts has been described before (Imayoshi *et al*., 2010; Imayoshi *et al*., 2008; Son *et al*., 2020), but to our knowledge, our study is the first to report corpus callosum defects upon alterations of Notch signaling in mice.

In NICD mice, we observed increased number of SATB2+ cells together with the presence of Probst bundles, indicating that callosal axons are present but unable to cross the midline. One possibility is that the defects in the corpus callosum may be directly caused by the changes in the cortical cell ratios observed in our different models. The pioneering axons –the first axons to cross the midline during development- are known to guide later axons, and experimental approaches indicate that in the absence of the pioneering axons, later axons are unable to find the correct path (Koester & O’Leary, 1994). SATB2+ neurons are normally detected from E13.5 as CUX1+ upper layer neurons, but a small percentage colocalize with CTIP2 or TBR1 (Alcamo *et al*., 2008). Thus, it is possible that the lack of CTIP2+ SATB2+ or TBR1+ SATB2+ pioneering axons is causing this phenotype. Alternatively, deficiencies in the midline glia (glial wedge and indusium griseum) which secrete guidance cues or changes in the expression of the appropriate receptors in the SATB2+ neurons could also result in the failure to cross the midline. In this direction, our RNAseq results indicate that several guidance receptors are altered upon NICD overexpression, including *Slit1, DCC1, UNC5a, Plxna4*, and *Nrp1* (Supplementary Table 1), even though our analyses were restricted to RGs and not postmitotic neurons.

In humans, agenesis and dysgenesis of the corpus callosum (ACC and DCC) are among the most frequent malformations in the brain with a reported incidence ranging between 0.5 and 70 in 10,000 births. These defects often correlate with mental retardation, hydrocephalus, visual problems, and autism (Schell-Apacik *et al*, 2008). Although in many patients the genetic cause of these disorders remains unclear, abnormal corpus callosum has been described in patients with pathogenic variants of the Notch ligand *DLL1* (Fischer-Zirnsak *et al*, 2019). We believe our animal lines offer a novel suitable model to investigate the pathophysiology of these human defects.

### Notch regulates radial glia cell cycle length and cortical neurogenesis

In the present study, we show that the switch between generating deep to upper-layer projection neurons is regulated by Notch, as RGs labeled with EdU generated upper-layer fates sooner in NICD mice and they generated deep-layer cells for longer periods upon the expression of dnMAML. These results parallel phenotypes observed in Hes5 KO and Hes5-overexpression models where the timing of neurogenesis was also affected (Bansod *et al*., 2017).

Since the length of the cell cycle is longer in NICD RGs, the fate acquisition changes do not seem to correlate with the number of cell divisions. However, the length of the cell cycle and the timing of cell cycle exit could be influencing these neurogenic fate decisions. For example, during bristle patterning in *Drosophila*, Notch signaling controls cell cycle progression such that cells with elevated Notch signaling divide first while those with lower signaling extend their G2 phase, making them more sensitive to lateral inhibition and consequently change their cell fate (Hunter *et al*, 2016). Our data could fit a similar model in which NICD extends RG cell cycle time, making these cells more susceptible to fate determinant factor(s).

Our data support the idea that fate decisions are decided at the progenitor stage. In this direction, pro-neural bHLH transcription factors, a family of transcriptional regulators known to play key roles in fate determination, are expressed in the terminal cell cycle of neural progenitors from S or G2 (Prasov & Glaser, 2012), and are known to regulate both cell cycle exit and fate choices (Castro *et al*, 2011; Maurer, 2018; Nguyen *et al*, 2006). Surprisingly, we also detected the presence of transcripts normally associated with specific subpopulations of postmitotic neurons in our sequencing experiments using purified RGs. As suggested before, low levels of these mRNAs in RGs may not lead to detectable protein expression but might prime the cells for differentiation upon cell cycle exit. Interestingly, while we distinguished significant differences in the expression of IP markers (*EOMES/Tbr2*) and deep layer genes (*Tbr1, Myt1, Ctip2*), we did not observe any differences in upper-layer markers (*Cux1, Satb2*). Published reports have shown that TBR1/CTIP2 and TBR1/FEZF2 initially overlap in their expression before genetic repression and depression networks establish the distinct layer subtype identities (Toma *et al*, 2014). For instance, FEZF2 is a transcriptional repressor that acts by repressing genes that would be inappropriate for layer V, including *Tbr1* (Srinivasan *et al*, 2012) and layer II-IV genes (Tsyporin *et al*, 2021). Our data may suggest that the reduction in deep layer genes in RGs may be affecting the expression of upper layer gens later (*i*.*e*., upon cell cycle exit).

### miRNAs downstream of Notch are required for upper-layer neuron fate acquisition

We identified two miRNA clusters –*miR100hg* and *miR99ahg*-with increased expression in NICD cortices. Both clusters encode for let-7, miR-125b, and miR-99a/100. Let-7 and miR-125b are essential regulators of developmental timing in various organisms (Ambros & Horvitz, 1984; Pasquinelli *et al*, 2000; Wu *et al*, 2012). In the mammalian retina, let-7 and miR-125b regulate the switch from progenitors that produce early cell fates to retinal progenitors that yield late-born cell fates (La Torre *et al*., 2013; Xia & Ahmad, 2016), while in the cortex, let-7 has been recognized as an important factor to maintain homeostasis (Fernandez *et al*, 2020) and vital in the generation of late cell types (Patterson *et al*., 2014; Shu *et al*., 2019). Moreover, we have recently shown that let-7 also regulates progenitor cell cycle dynamics in the cortex (Fairchild *et al*, 2019). While there is limited literature on the roles of miR-99a/100 in the CNS, the miR-99a/100, let-7, miR-125b tricistrons have been shown to regulate hematopoietic progenitor homeostasis (Emmrich *et al*, 2014).

Strikingly, the inhibition of these miRNAs activity using specific miRNA sponges is sufficient to partially rescue the NICD phenotype. We have previously shown that let-7 inhibition leads to a shortening of the S/G2 phase of cell cycle, but further inquiry is needed to discern if the effects of these sponges are mediated by the regulation of cell cycle or through other downstream targets. A widely-recognized target of let-7 is the chromatin remodeler HMGA2 (Lee & Dutta, 2007), a known important regulator of neocortical neurogenesis (Shu *et al*., 2019) but let-7 also regulates the levels of the nuclear receptor TLX (Zhao *et al*, 2010) and the cell cycle genes Cyclin D1, Cyclin D2, CDK4, CDK6 and CDC25A(Bueno & Malumbres, 2011). Further experiments aimed at understanding the molecular mechanisms downstream of let-7 and the possible cooperative activities between the different miRNAs will shed light on the machinery that instructs cortical fate acquisition.

Importantly, *miR100hg* is located in a human chromosome region (11q24.1) whose deletion is associated with Jacobsen syndrome (JBS, OMIM #147791). This syndrome involves intellectual disability, abnormal head shape, microphthalmia, and increased likelihood of autism spectrum disorders (Akshoomoff *et al*, 2015; Favier *et al*, 2015; Guerin *et al*, 2012). The JBS patient deletions range from 7 to 20 Mb. The heterogeneity of phenotypes supports the hypothesis that JBS is a contiguous gene deletion syndrome where the loss of different combinations of genes causes particular phenotypes. Owing to the rarity of reported cases reported, it has not yet been possible to tease out the requirements of individual genes, including miR100hg. Notably, a rare microtriplication of 1.8 Mb in the 11q24.1 region (partial trisomy), which includes *miR100hg*, and only a handful of other genes (11 in total), also results in intellectual disability with severe verbal impairment (Beneteau *et al*, 2011). Thus, a further understanding of the roles that *miR99ahg* and *miR100hg* play in telencephalic development will also allow us to gain further insights into the molecular underpinnings of developmental brain disorders.

## ACKNOWLEDGMENTS

We want to thank all members of the Brown, Simó, and La Torre laboratories for their helpful insights. We also want to thank Dr. Tom Glaser and Dr. Nick Marsh-Armstrong for their valuable comments and generosity with reagents. This study was supported by the National Institute of Neurological Disorders and Stroke (National Institutes of Health) [R21 NS101450 to S.S. and A.L.T., R01 NS109176 to S.S.] and National Eye Institute [R01 EY013612 and R01 EY031724 to N.L.B]. We also benefited from the use of the National Eye Institute Core Facilities [supported by P30 EY012576] and the University of California Davis Flow Cytometry Shared Resource Laboratory with funding from the NCI [P30 CA0933730] and NCRR [C06-RR12088, S10 RR12964] with technical assistance from Ms. Bridget McLaughlin and Mr. Jonathan Van Dyke. The sequencing library preparations and the sequencing were carried out at the UC Davis Genome Center DNA Technologies and Expression Analysis Core, supported by NIH Shared Instrumentation Grant 1S10OD010786-01 and analyzed by the UC Davis Bioinformatics Core. Graphical abstract was created with BioRender.

## EXPERIMENTAL MODELS AND SUBJECT DETAILS

### Animals

All animals were used with approval from the University of California Davis Institutional Animal Care and Use Committees and housed and cared for in accordance with the guidelines provided by the National Institutes of Health. B6N.129-*Gt(ROSA)26Sor*^*tm1(MAML1)Wsp*^/J (ROSA26^loxP-stop-loxP-dnMAML1^) and *Notch1*^*tm2Rko*^/GridJ (Notch1^f/f^) were generous gifts from Dr. Ivan Maillard and Dr. Raphael Kopan, respectively. *Gt(ROSA)26Sor*^*tm1(Notch1)Dam*^/J (ROSA26^loxP-stop-loxP-Notch1-ICD^) and B6.129S2-*Emx1*^*tm1(cre)Krj*^/J (Emx1-Cre) mice were obtained from The Jackson Laboratory. All animals are currently available at The Jackson Laboratory (Cat. #008159 (Murtaugh *et al*., 2003), #032613 (Tu *et al*., 2005), #006951 (Yang *et al*., 2004) and #005628 (Gorski *et al*., 2002), respectively). To drive NICD and dnMAML expression in the developing mouse telencephalon, ROSA26^loxP-stop-loxP-Notch1-ICD^ or ROSA26^loxP-stop-loxP-dnMAML1^ were crossed with Emx1-Cre/+ mice. To generate Emx1-Cre/+; Notch1^f/f^ (Notch1cKO) mice, Notch1^f/f^ were crossed with Emx1-Cre*/+* mice to generate an intermediate stock and then Emx1-Cre/+; Notch1^f/+^ were bred with Notch1^f/f^.

### Constructs

The MSCV puro let-7 sponge was a gift from Dr. Phil Sharp (Addgene plasmid #29766) (Kumar *et al*, 2008), the pRNA-U6-let-7 sponge was a gift from Dr. Phillip Zamore (Addgene plasmid #35664), and the MG-miR-125b-sponge-bulge was a gift from Dr. David Baltimore (Addgene plasmid #45790) (Chaudhuri *et al*, 2012). The miR-100 sponge was designed using the miRNAsong algorithm (Barta *et al*., 2016) (5’-CACAAGTTCGGATCTACGGGTTAATTCACAAGTTCGGATCTACGGGTT-3’). MiR-100 sponge sequences were cloned in tandem into a pCAG backbone(Simo & Cooper, 2013) to obtain a 12mer miR-100 sponge sequences. miR-100 sponge is expected to also target miR-99a based on sequence homology.

### Histology and Immunohistochemistry

Brains were collected at indicated ages and fixed in 3.7% formalin/PBS by submersion overnight at 4ºC. Tissues were cryoprotected with 30% sucrose/PBS solution. Next, brains were embedded in Optimum Cutting Temperature (OCT) compound (Tissue-Tek, Torrance, CA) and quickly frozen using dry-ice. OCT embedded brain blocks were cryo-sectioned on a coronal plane (15 μm). Immunostainings were performed in free-floating sections with agitation. Sections were blocked with PBS, 0.3% Triton X-100 and 5% milk or 10% normal donkey serum for 1h at room temperature. Blocking solution was used for primary antibody incubation (overnight, 4°C). After primary antibody incubation, free-floating sections were washed three times in PBS/0.1% Triton X-100 (10 min each). Species-specific, fluorescently-labelled secondary antibodies were used in blocking solution (90 minutes, room temperature). 4’,6-diamidino-2-phenylindole (DAPI) (Sigma-Aldrich) was used for nuclear staining. The list and concentrations of antibodies used in this work are described in the table below. Images were taken in a Fluoview FV3000 confocal microscope (Olympus, Center Valley, PA) or Axio Imager.M2 with Apotome.2 microscope system (Zeiss, Dublin, CA). All images were assembled using Photoshop and Illustrator (Adobe, San, Jose, CA).

| **Antibody** | **Source** | **Catalog** | **Concentration** |
| --- | --- | --- | --- |
| Anti-BrdU (mouse) | ThermoFisher | #B35128 | 1:100 |
| Anti-Calbindin (Mouse) | Millipore Sigma | #C9848-100UL | 1:200 |
| Anti-Cleaved Caspase-3 (Rabbit) | Cell Signaling Technologies | #9664S | 1:200 |
| Anti-CTIP2 (Rat) | Abcam | #ab18465 | 1:500 |
| Anti-CUX1 (Rabbit) | Proteintech | #11733-1-AP | 1:1000 |
| Anti-CUX1 (Rabbit) | Santa Cruz Biotechnologies | #sc-13024,<br>discontinued | 1:50 |
| Anit-DKK3 (Rabbit) | Sino Biological | #50247-RP02 | 1:200 |
| Anti-FOXG1 (Rabbit) | Abcam | #ab196868 | 1:100 |
| Anti-GFAP (Mouse) | Antibodies Incorporated | #73-240 | 1:100 |
| Anti-L1 (Rabbit) | Millipore Sigma | #MAB5272 | 1:500 |
| Anti-MSX1 (Goat) | R&D Systems | #AF5045; | 1:100 |
| Anti-PAX6 (Rabbit) | BioLegend | #901301; | 1:250 |
| Anti-PH3 | ThermoFisher | #PA5-17869 | 1:100 |
| Anti-Reelin (Mouse) | Millipore Sigma | #MAB5364; | 1:100 |
| Anti-RFP (Rabbit) | Novus Biologicals | #NBP2-25157; | 1:1000 |
| Anti-RFP (Goat) | Origene | #AB0040-200; | 1:1000 |
| Anti-SATB2 (Mouse) | Abcam | #51502; | 1:200 |
| Anti-TBR1 (Rabbit) | Invitrogen | #PA5-34582; | 1:1000 |
| Anti-TBR2 (Rabbit) | Abcam | #ab23345; | 1:200 |
| Anti-β-III-Tubulin (TUJ1) (Mouse) | BioLegend | #801201; | 1:500 |

### *In utero* electroporation

*In utero* microinjection and electroporation was performed at embryonic day (E)13 as described previously (Simo & Cooper, 2013), using timed pregnant NICD mice. For control electroporations, DNA solutions containing 0.5 mg/ml pCAG-ChFP plasmids were mixed in 10 mM Tris, pH 8.0, with 0.01% Fast Green and 1 μl of the solution was injected per embryo. Tweezertrodes electrodes (BTX) with 5-mm pads were used for electroporation (five 50ms pulses of 30V). To express the miRNA sponges, a solution containing 1 mg/ml of each sponge individually, or combined, and 0.5 mg/ml pCAG-ChFP was used. All experimental manipulations were performed in accordance with protocols approved by the University of California, Davis Institutional Animal Care and Use Committee (IACUC). At P0, electroporated brains were collected and processed as described.

### FlashTag NSC labelling and FACS

Labelling of cortical neuronal progenitors with carboxyfluorescein esters (CFSEs) was achieved as described elsewhere (Govindan *et al*., 2018). Briefly, 1μl of a 5mM solution of CellTrace CFSE (from the CellTrace CFSE Cell Proliferation Kit, Invitrogen #C34554) and 0.01% FastGreen in DMSO was injected into the 3^rd^ ventricle of E13.5 control and NICD embryos. Dams were allowed to recover and injected embryos were collected 1 hour post-injection. Embryonic cortices were dissected individually and dissociated into single cells using Papain Dissociation System (Worthington, # LK003150) using manufacturer’s protocol. Cells were resuspended in FACS media (DMEM/F12 without phenol red supplemented with 10% FBS and B-27) and sorted using a Beckman Coulter Astrios EQ Cell Sorter.

### RNA and miRNA sequencing

Total RNA from control (n=3) and NICD (n=5) sorted cells was extracted using the Total RNA Purification Plus Kit (NORGEN Biotek Corp., #48300). Gene expression profiling was carried out using a 3’-Tag-RNA-Seq protocol. Barcoded sequencing libraries were prepared using the QuantSeq FWD kit (Lexogen, Vienna, Austria) for multiplexed sequencing according to the recommendations of the manufacturer using both, the UDI-adapter and UMI Second-Strand Synthesis modules (Lexogen). The fragment size distribution of the libraries was verified via micro-capillary gel electrophoresis on a LabChip GX system (PerkinElmer). Barcoded miRNA-Seq libraries were prepared using the NEXTflex Small RNA Sequencing kit V3 (PerkinElmer) with sequence randomized adapters according to the recommendations of the manufacturer. The fragment size distribution of the libraries was verified via micro-capillary gel electrophoresis on a Bioanalyzer 2100 (Agilent). Both sets of libraries were quantified by fluorometry on a Qubit instrument (Life Technologies, Carlsbad, CA), and pooled in equimolar ratios. The library pools were quantified by qPCR with a Kapa Library Quant kit (Kapa Biosystems/Roche). Finally, both library sets were sequenced on a HiSeq 4000 sequencer (Illumina) with single-end 100 bp reads.

### EdU and BrdU labelling

For EdU birth-dating experiments, pregnant dams were injected intraperitoneally with 25mg EdU/kg body weight at E13.5 and pups were sacrificed at birth. For the EdU/BrdU dual window labeling experiment, pregnant dams were injected intraperitoneally with 12.5mg/kg body weight of EdU at E13.5 and injected again with 12.5mg/kg body weight of BrdU after 2 hours. Brains were collected 30 minutes post BrdU injection. EdU was detected following manufacturer instructions (ThermoFisher Scientific, #C10337). Prior to detection of BrdU by immunofluorescence as previously described, tissue was treated with hot 10mM sodium citrate pH 8 for 20 min followed by an acid wash (2N HCl and 0.5% Triton X-100 in PBS) for 1 hour at room temperature.

### Statistical Methods

Specific number of biological replicates (Ns) and statistical methods used are specified in each figure or figure legend. For cortical thickness, the thickness of the somatosensory cortex in three consecutive brain slices was measured and averaged per brain. The same strategy was used to measure the corpus callosum thickness. The hippocampal area was measured from both hippocampal hemispheres of three sections containing the dorsal hippocampus in which the habenula was visible. These measures were averaged and represented the results for a single brain. For histological and IUE quantifications, single or double fluorescently labelled cells were quantified for at least three consecutive sections in each brain and their results averaged.

To measure cell distribution in the cortex we used RapID (Sekar *et al*., 2021). Briefly, a grid containing eight equally-sized bins was manually placed in the cortex with bin 1 mainly covering the marginal zone and bin 8 covering the intermediate zone. The quantification of fluorescently labelled cells was automatically determined by the software.

All statistical analyses and plot generation were performed using Prism 9 (GraphPad, San Diego, CA).

## SUPPLEMENTARY FIGURE LEGENDS

**Supplementary Figure 1. NICD and dnMAML mice**

**A**. At P14, NICD mice are smaller in size compared to their control littermates. **B**. At P0, the size of the brain is significantly smaller in dnMAML mice compared to their littermate controls but no significant differences were observed in NICD. Scale bar B-D: 5mm.

**Supplementary Figure 2: Agenesis of the corpus callosum in NICD mice**

**A-B**. At P0, NICD cortices exhibit increased ratios of SATB2+ callosal neurons (green) compared to their littermates, DAPI (blue) was used for counterstaining. P-value was obtained using Student’s T-test. Scale bar: 100 microns. **C**. Upon in utero electroporation of mCherry (red), labeled axons (while arrows) cross the midline in control animals but result in aberrant bundles in NICD mice. Scale bars: 100μm A-B; 250μm C.

**Supplementary Figure 3: Hippocampal defects in NICD and dnMAML mice**

**A**. Caspase3 (green) and GFAP (red) immunolabeling of control and NICD hippocampi at P0, counterstained with DAPI (blue). Scale bar: 100 microns. **B**. Caspase3 (green) immunolabeling of control and NICD neocortices at P0. White arrows point at Caspase3+ cells. Scale bar: 70 microns. **C**. Quantification of Caspase3+ cells in hippocampus and cortex, p-values were obtained using Student’s T-test. **D**. Immunolabeling using DKK3 (red) and Calbindin (green) antibodies, counter-stained with DAPI (blue) of dnMAML hippocampal section. Note the ectopic location of some DKK3+ cells (white arrows). Scale bars: 100μm A, 70μm B, 200μm D.

**Supplementary Figure 4: Aberrant production of Cajal-Retzius cells in NICD and dnMAML mice**

**A**.NICD cortices exhibit less Reelin+ (white label, yellow arrows) than their littermate controls. Scale bar: 50 microns. MZ: marginal zone. **B**. Immunolabeling with Reelin (red) in control and NICD hippocampal sections. The white inset box is shown in E. LV: lateral ventricle, DG: dentate gyrus. Scale bar: 200 microns. **C-D**. Quantification of number of Reelin+ cells/area in NICD and dnMAML hippocampi. P-Values were obtained using Student’s T-tests. **E**. Only a handful of Reelin+ cells (white, noted with yellow arrows) are detected in NICD hippocampi. Scale bars: 50 μm A, 200μm B, 100μm E

**Supplementary Figure 5: Lamination defects in dnMAML neocortices**

Brain coronal sections immunolabeled against CTIP2 (green), TBR1 (red), and counterstained with DAPI (blue). Scale bar: 500μm.

**Supplementary Figure 6: NotchcKO phenotypes**

**A-B**. Brain coronal sections immunolabeled against CTIP2 (green), TBR1 (red), and counterstained with DAPI (blue). The thickness of the corpus callosum is indicated with a white bar. Scale bar: 500 microns. CC: corpus callosum; LV: lateral ventricle. C-D. Hippocampal sections stained against Calbindin (green) and DKK3 (red) and counterstained with DAPI (blue). Scale bar: 250 microns. CA1-CA3: Cornu Ammonis hippocampal regions, DG: dentate gyrus. **E-F**. Cortical sections stained with TBR1 (green) and CTIP2 (magenta), and counterstained with DAPI (blue). Scale bar: 100 microns. **G-H:** Quantifications of CTIP2+ and TBR1+ cells/area, respectively in Control and NotchcKO samples. **I-J**. Quantification of ratio of EdU+ cells colabeled with either CTIP2 or TBR1. For all quantifications (G-J), p-values were obtained using Student’s T-tests. Scale bars: 500μm A-B, 250μm C-D, 100μm E-F.

**Supplementary Figure 7: RNA sequencing**

**A**. Multidimensional scaling analysis (MDS) showing control samples (orange) and NICD samples (teal). **B**. Gene Ontology (GO) analyses using PANTHER Classification System.

**Supplementary Table 1. RNA-seq Differential Expression**

Normalized counts of NICD vs Control samples.

**Supplementary Table 2. microRNA-seq Differential Expression**

Normalized counts of NICD vs Control samples.

