## Supplementary figures and images for "Notch directs telencephalic development and controls neocortical neuron fate determination by regulating microRNA levels"

### Supplementary Figures

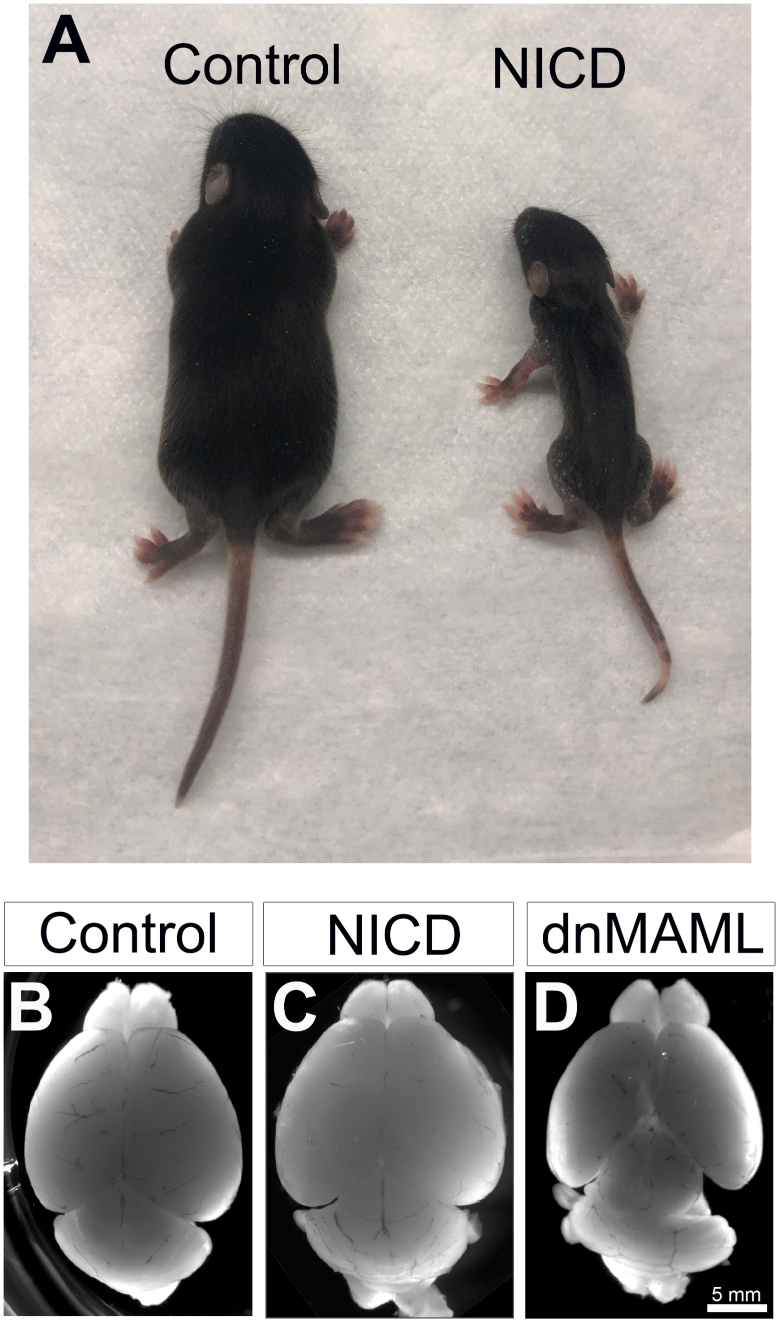

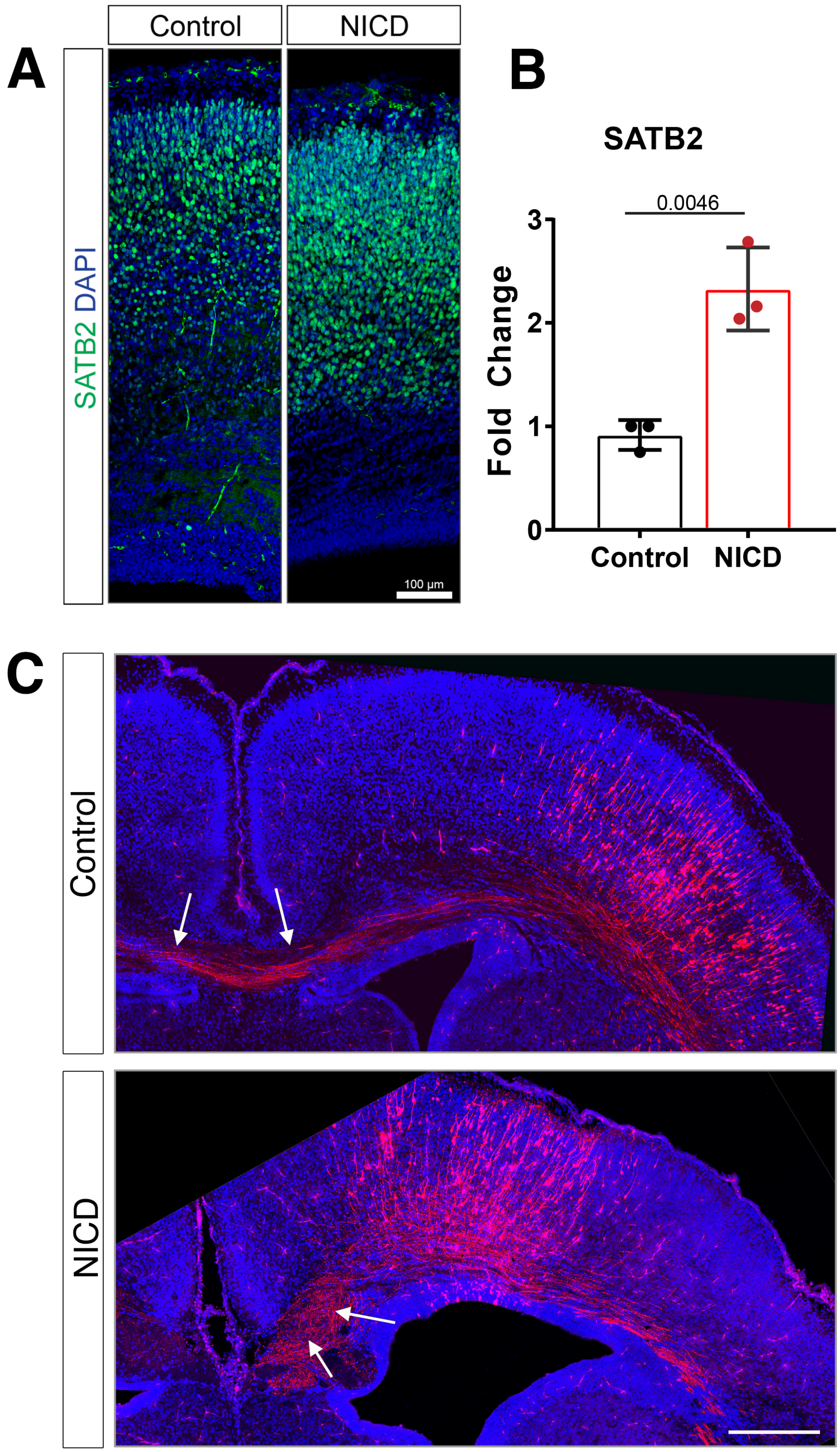

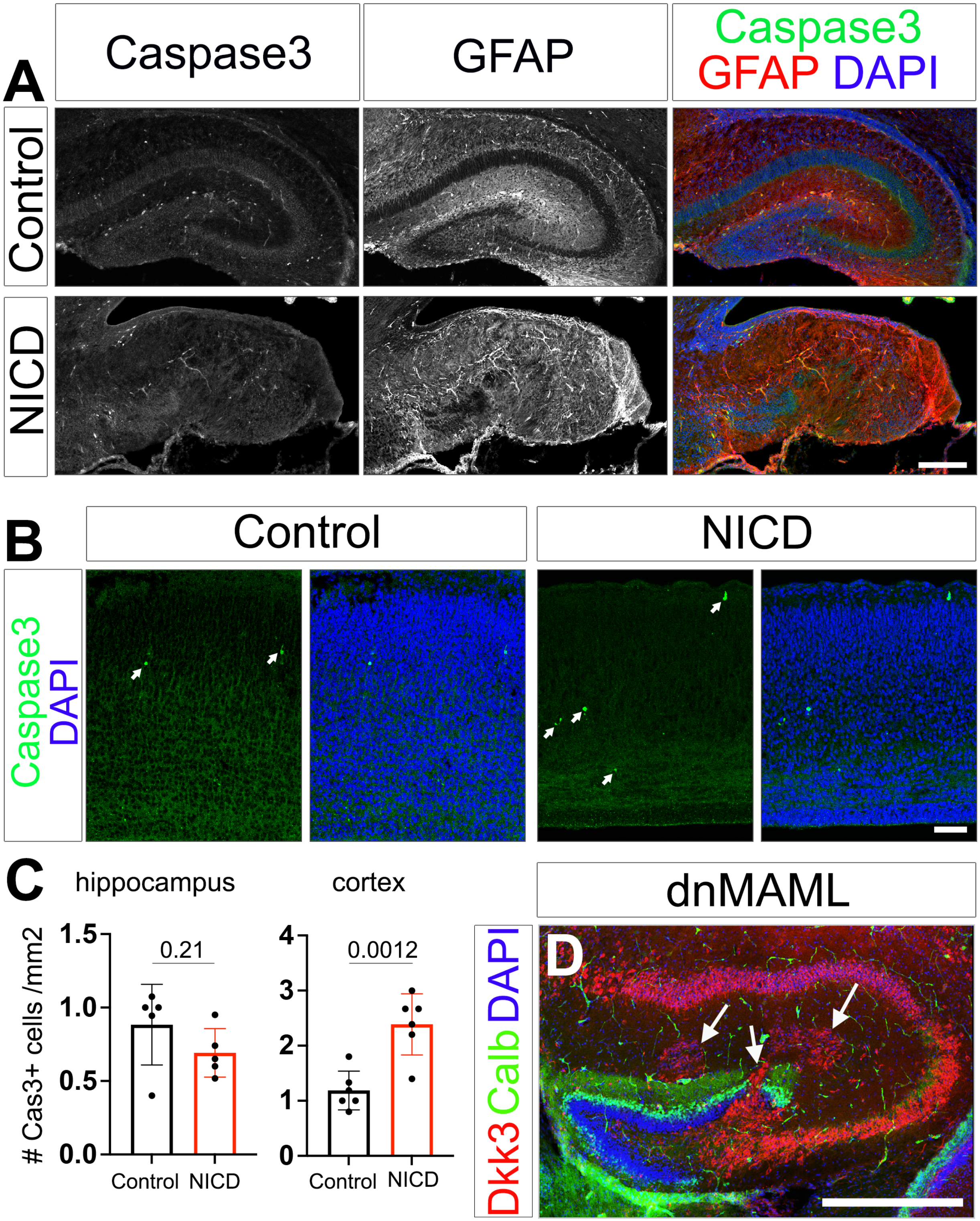

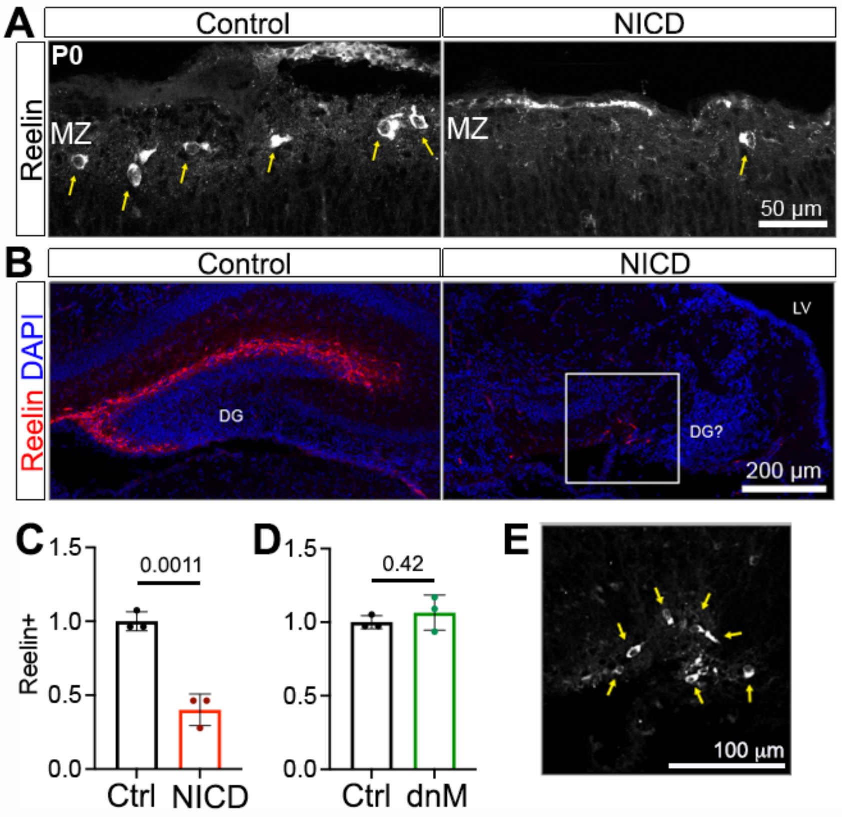

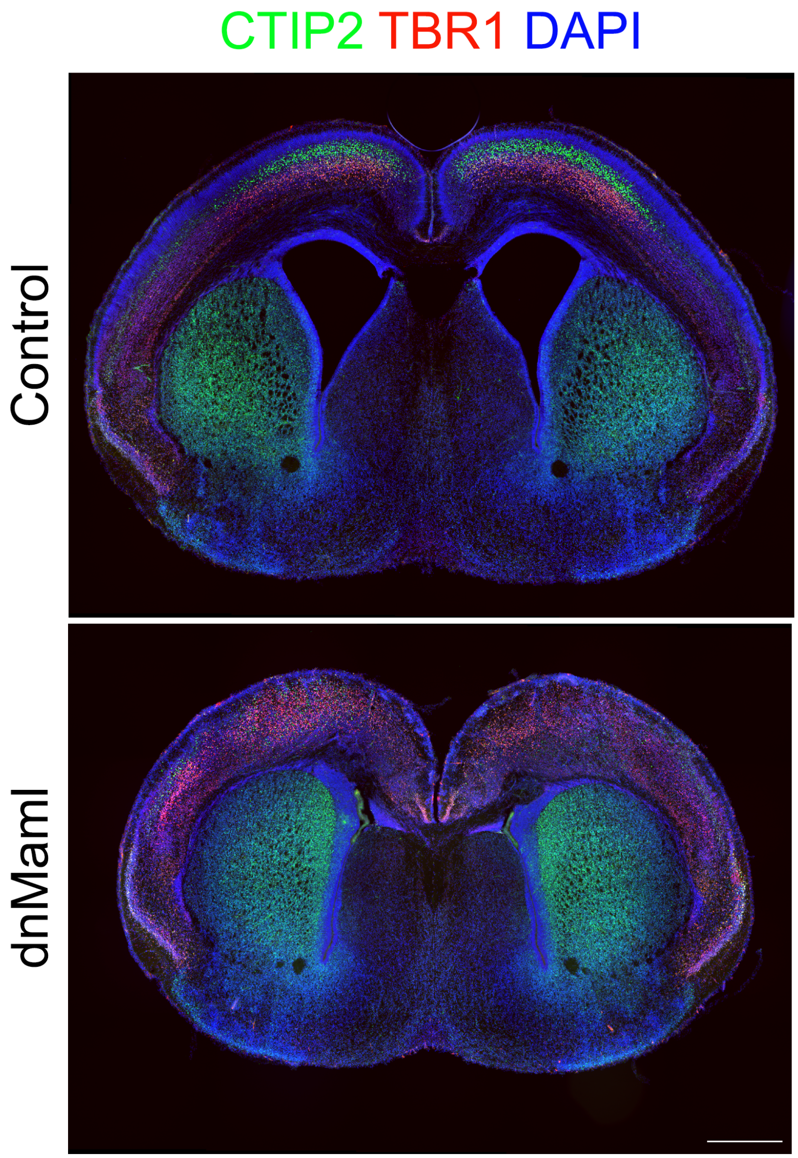

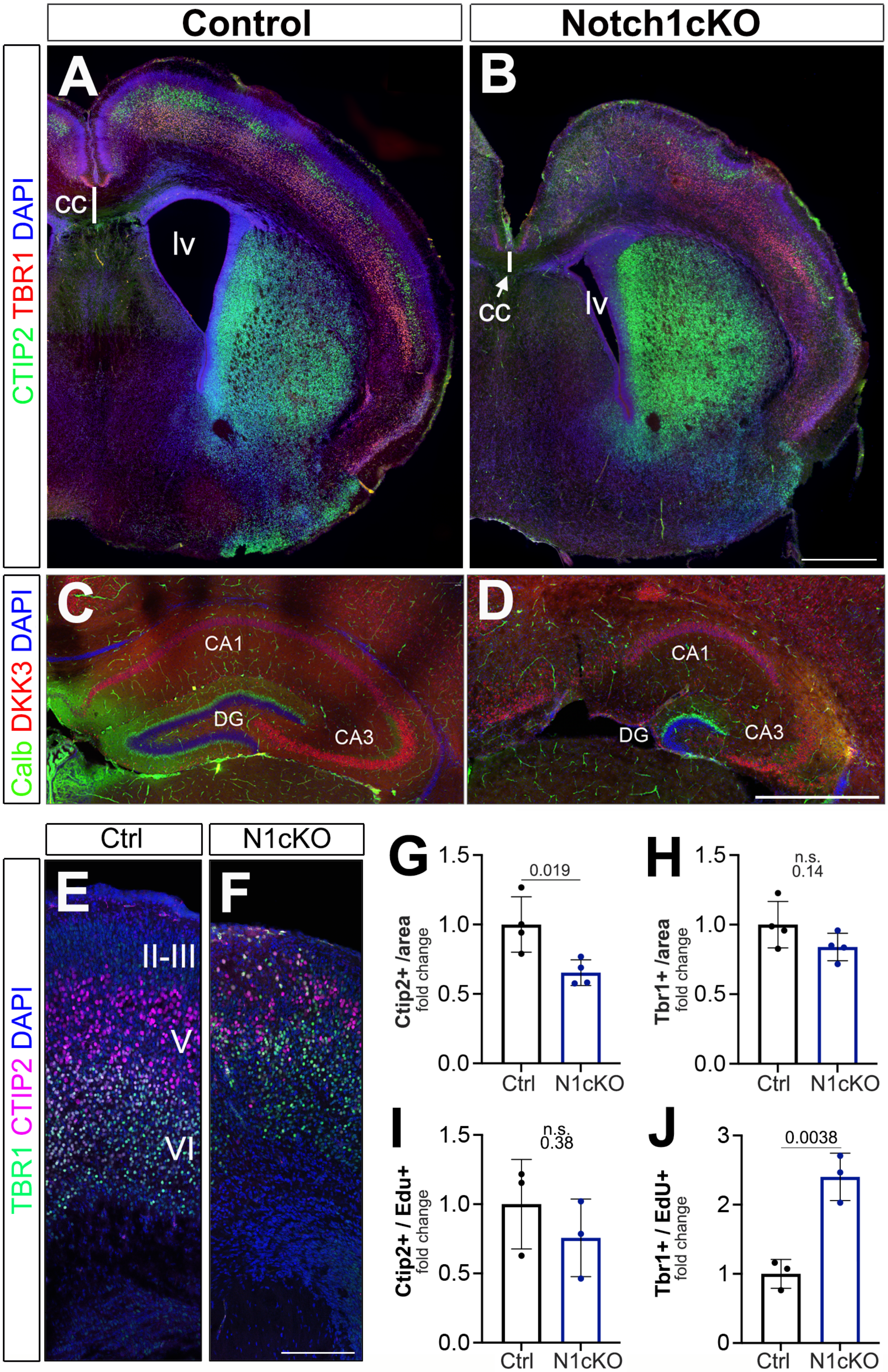

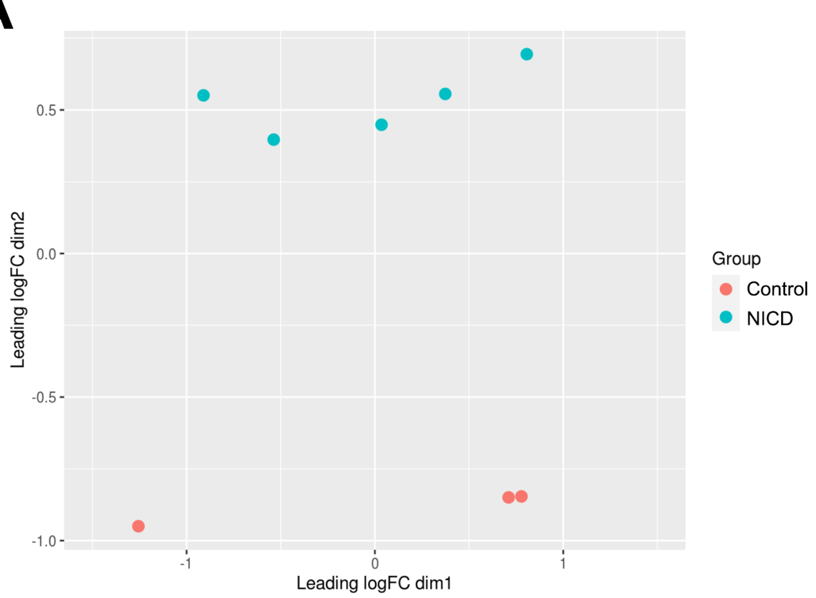

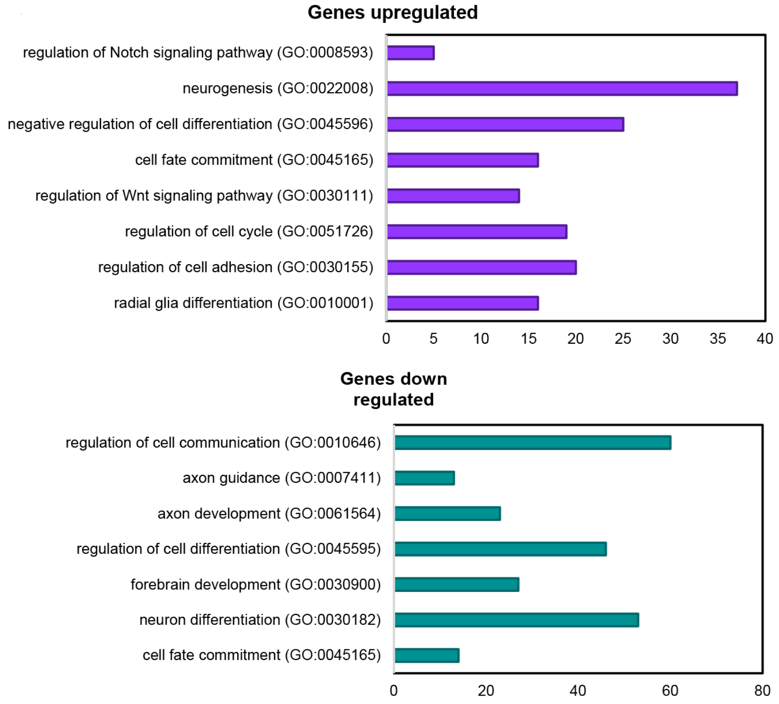
